# Spatial heterogeneity in carbohydrates and their utilisation by microbes in the high North Atlantic

**DOI:** 10.1101/2023.05.11.540373

**Authors:** Taylor Priest, Silvia Vidal-Melgosa, Jan-Hendrik Hehemann, Rudolf Amann, Bernhard M. Fuchs

**Author notes:** **Corresponding author** Taylor Priest.

## Abstract

Carbohydrates are chemically and structurally diverse, represent a substantial fraction of marine organic matter and are key substrates for heterotrophic microbes. Studies on carbohydrate utilisation by marine microbes have been centred on phytoplankton blooms in temperate regions, while far less is known from high-latitude waters and during later seasonal stages. Here, we combine glycan microarrays and analytical chromatography with metagenomics and metatranscriptomics to show the spatial heterogeneity in glycan distribution and their utilisation by microbes in Atlantic waters of the Arctic during late summer. The composition and abundance of monomers and glycan structures in POM varied with location and depth. Complex fucose-containing sulfated polysaccharides, known to accumulate in the ocean, were consistently detected, suggesting limited degradation by microbes. In contrast, the more labile β-1,3-glucan exhibited a patchy distribution, indicating local variations in primary productivity and rapid utilisation. Metatranscriptomics showed active and dynamic microbial populations that targeted specific glycans. Gene transcription of carbohydrate-active enzymes revealed narrow substrate niches for specialists, involving compounds such as α-mannans and alginate, along with the targeting of communal substrates, such as laminarin, by multiple populations. The observed spatial heterogeneity indicates that local biological and physical processes continue to shape the carbohydrate pool during late summer in high latitude waters and microbial populations are active and responsive to such changes.

## INTRODUCTION

Marine carbohydrates are chemically and structurally diverse, and represent a substantial fraction of characterised organic matter [1]. The diversity of glycans, which are the polymeric carbohydrates, emerges from the alternative linkage types, alpha and beta, of carbon atoms between more than ten available monomers along with substitutions by a range of other chemical groups [2]. Micro- and macro-algae are the primary synthesisers of glycans in the ocean, wherein they serve structural, storage and protective functions. Glycans can constitute between 13 – 90% of algal carbon [3]. Single-celled algae, or phytoplankton-derived glycans range from low-molecular weight (LMW) oligosaccharides to complex high-molecular weight (HMW) polysaccharides, with varying composition across taxa, life cycle stage and environmental conditions [4, 5]. Through exudation, cell death and lysis, various glycans are released and integrated into the particulate and dissolved organic matter pools (POM and DOM) [6–8], which are separated based on the size of particles. Once released or outside the cell, glycans can become energy for heterotrophic microbes.

Carbohydrate utilisation is common in bacteria and archaea, but the mechanisms employed and the degradative capabilities vary [9–11]. Many species take up mono-, di- and trisaccharides into the cell through porins or transporters, while longer oligosaccharides require specialised systems, such as TonB-dependent transporters (TBDTs) or other outer membrane proteins. Usually these have a high specificity to discrete glycan structures [12]. For polysaccharides, microbes must first depolymerize the structure extracellularly with excreted or outer membrane-bound glycoside hydrolases (GHs) or polysaccharide lyases (PLs), followed by uptake of the oligosaccharides and subsequent cleavage in the periplasm [12, 13]. These enzyme classes, together with carbohydrate-binding modules (CBMs) and carbohydrate esterases (CEs), are collectively referred to as carbohydrate-active enzymes (CAZymes). CAZymes are classified into families based on protein sequence similarity, with each family also containing at least one biochemically characterized protein [14, 15]. Many families are monospecific to certain glycosidic linkage types within polysaccharides while others are divided into sub-families based on specificity of target linkages [16]. Unlike those from land plants, algal glycans are decorated with sulfate esters, which require sulfatase enzymes for complete degradation. The CAZyme, sulfatase and transporter gene profiles thus acts as the blueprint for the glycan degradation potential of microbes [10, 17].

Carbohydrate utilisation by microbial populations exhibit spatial and temporal variations. The rate of hydrolysis and the substrate spectrum of extracellular CAZymes decreases and narrows with depth [18] and with distance to the coast [19]. In addition, a broader spectrum of CAZyme activities is measureable in temperate compared to high-latitude waters, indicating latitudinal gradients [20]. Temporal shifts in CAZyme, sulfatase and transporter gene profiles are also evident following spring phytoplankton blooms [21]. These patterns are congruent with dynamic changes in microbial community composition. In particular, community-level patterns are shaped by the presence and composition of specialised carbohydrate degraders, such as *Bacteroidetes. Bacteroidetes* typically harbour large CAZyme repertoires [10, 17] and exhibit successional dynamics following spring phytoplankton blooms [21]. These dynamics indicate glycan-based niche partitioning [22, 23]. Detailed assessments on microbial carbohydrate utilisation have been focused on spring phytoplankton blooms in temperate ecosystems. Whether similar patterns occur at later seasonal stages and in high-latitude waters remains unknown.

In this study, we combine analytical techniques with meta’omics approaches to explore the distribution of glycans and the dynamics of glycan utilisation by microbes in Atlantic waters of the Arctic during late summer. Combining glycan microarrays and high-performance anion-exchange chromatography provided insights into the distribution of monosaccharides and specific glycan epitopes in the upper euphotic zone. Concurrently, the application of PacBio HiFi read metagenomics and short-read metatranscriptomics allowed for the active fraction of the microbiome to be distinguished and the identification of population- and community-level glycan utilisation patterns. The reported data reveals diverse glycans with spatially heterogeneous distribution patterns along with active microbial populations targeting specialised and communal substrates.

## METHODS

### Sample collection

Seawater samples were collected from ten stations located in the eastern Fram Strait and around the Svalbard archipelago in September 2020 during the MSM95 Maria S. Merian research cruise [24]. A map of the sampling locations (Figure 1) was generated using publically available bathymetric data from the International Bathymetric Chart of the Arctic Ocean (IBCAO) [25, 26] and the QGIS v3.14.16-Pi [27] software. Seawater was collected using a CTD-rosette sampler from surface water (SRF), typically 2 m depth, and the bottom of the surface mixed layer (BML). The BML depth was defined by the beginning of the thermo- and halocline and drop in surface fluorescence values (Figure 1). One location was sampled twice over a two day period, with additional samples collected at 100 and 200 m depth, these are labelled as S1 and S6 (Supplementary Table S1). Of the water collected, 4 L was filtered sequentially through a 3 and 0.2 µm pore-size polycarbonate membrane filter (142 mm diameter) and immediately stored at −80 °C for ‘omics analysis. A second 4 L of seawater was filtered through a 0.7 µm pre-combusted Whatman Grade GF/F filter (47 mm diameter) and immediately stored at −80 °C for carbohydrate analysis.

**Figure 1.**
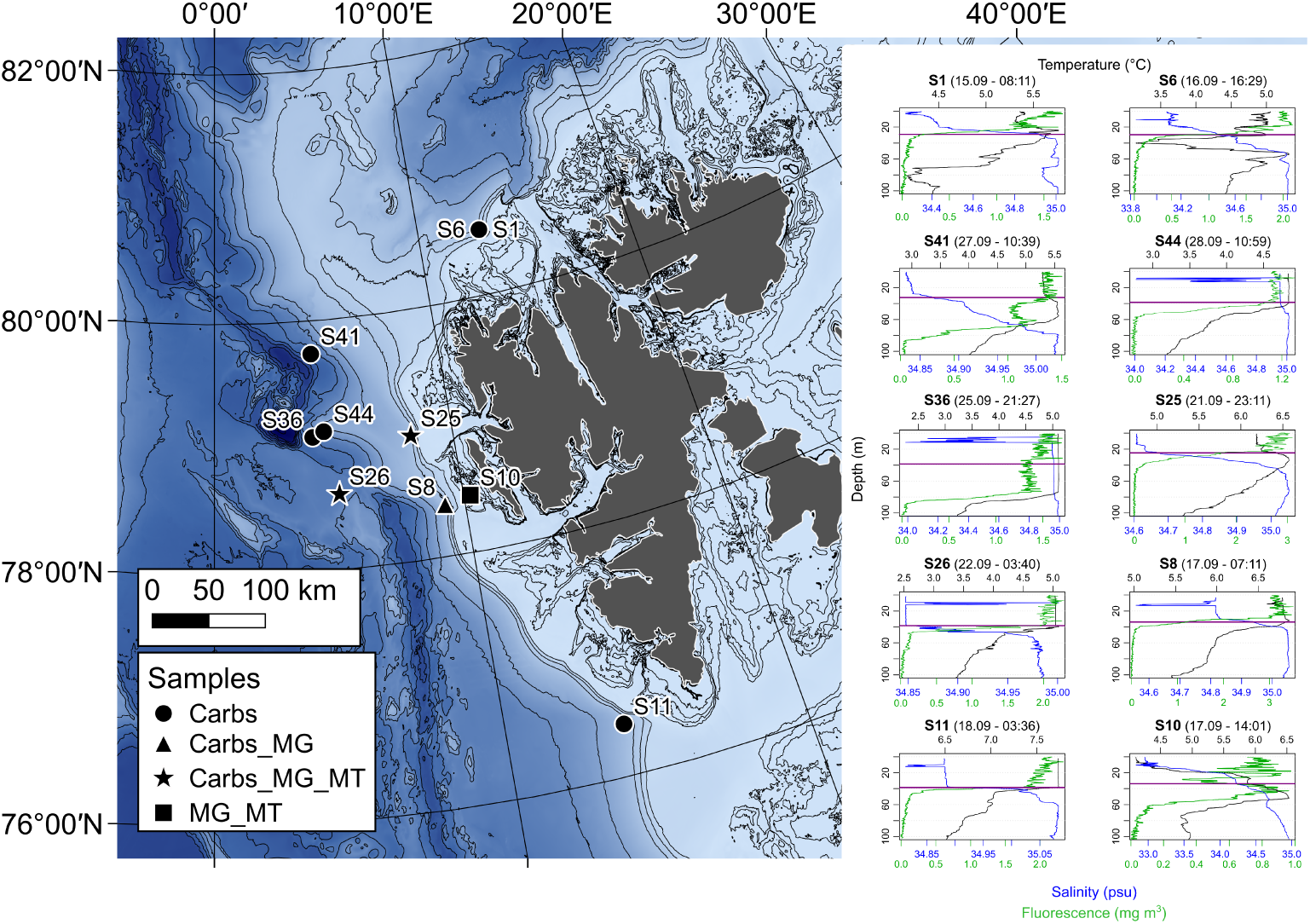
Bathymetric map with sampling locations, types of samples collected and vertical profiles from CTD casts. Map legend: Carbs = samples collected for carbohydrate analysis on particulate organic matter fraction, MG = sampled for metagenomics, MT = sampled for metatranscriptomics. Vertical profiles of temperature, salinity and fluorescence were derived from sensor measurements during CTD casts. The purple horizontal lines on each profile represent the bottom mixed layer (BML) sampling depth.

### Monosaccharide and polysaccharide analysis

The GF/F filters from all samples (ten stations and two depths) were cut into ten equally-sized circular pieces, with a diameter of 10 mm. The monosaccharide composition of and polysaccharide structures present on the filter pieces was analysed as described previously [28] and detailed in Supplementary Methods. First, two of the filter pieces were hydrolysed using acid (1 M HCI for 24 h at 100 °C) and the resulting monosaccharides were analysed using High-Performance Anion Exchange Chromatography with Pulsed Amperometric Detection along with monosaccharide standards. Acidic monosaccharides could not be detected due to a problem with the detector. The remaining filter pieces were subject to polysaccharide extraction using a sequential solvent protocol, with 1) MilliQ water, 2) 0.2 M EDTA (pH 7.5) and 3) 4 M NaOH with 0.1% w/v NaBH_4_. The identification and semi-quantitative analysis of polysaccharide compounds was performed using a microarray and antibody-based approach. Polysaccharide extracts were printed in quadruplicates onto nitrocellulose membranes (0.45 µm pore-size) using a microarray robot (Sprint, Arrayjet, Roslin, UK). The printed arrays were blocked for 1 h in 1 x PBS with 5% (w/v) non-fat milk powder (MPBS), followed by incubation for 2 h with polysaccharide-specific monoclonal antibodies (Supplementary Table S2). After incubation, the arrays were washed in PBS and incubated for 2 h in anti-rat, anti-mouse or anti-His tag secondary antibodies conjugated to alkaline phosphatase. Arrays were thoroughly washed in PBS and deionized water before being developed in a solution containing 5-bromo-4-chloro-3-indolylphosphate and nitro blue tetrazolium in alkaline phosphatase buffer (100 mM NaCl, 5 mM MgCl_2_, 100 mM Tris-HCl, pH 9.5). Developed arrays were scanned and the binding of each probe against each spotted sample was quantified using Array-Pro Analyzer 6.3 (Media Cybernetics). Mean antibody binding signal intensity (from four replicates) was determined for each extract of each sample, with the highest mean signal of the dataset being set to 100 and the remainder normalised accordingly.

### Metagenome and metatranscriptome sequencing

Filtered seawater samples of the 0.2 – 3 µm fraction from SRF and BML depths of four different stations (S8, S10, S25, S26) were subject to a dual nucleic acid isolation protocol using the DNA/RNA Mini Prep Plus kit from Zymo Research (Irvine, CA, USA), according to the manufacturer’s instructions. The quality of extracted DNA was assessed using capillary electrophoresis with a FEMTOpulse (Agilent), whilst RNA quality was assessed using a PicoChip on a Bioanalyser (Agilent, CA, USA). Ultra-low DNA libraries were prepared from the eight samples without further fragmentation by the protocol “Procedure & Checklist - Preparing HiFi SMRTbell® Libraries from Ultra-Low DNA Input” of PacBio (CA, USA). Libraries were sequenced on 4 x 8M SMRT cells on a Sequel II platform for 30 h with sequencing chemistry 2.0 and binding kit 2.0 (two samples multiplexed per SMRT cell). Four of the samples were additionally selected for metatranscriptome sequencing (S10_surface, S25_Surface, S25_BML, S26_BML). Illumina-compatible libraries were produced from extract RNA using the Universal Prokaryotic RNA-Seq library preparation kit, incl. Prokaryotic AnyDeplete® (Tecan Genomics, CA, USA). Libraries were sequenced on a HiSeq 3000 platform with 2 x 150 bp paired-end read mode.

### HiFi read taxonomic classification

A custom pipeline was employed to taxonomically classify metagenomic HiFi reads against a GTDB-based protein database. A Diamond blast (v0.9.14) [29] database was generated from the gene amino acid sequences of all GTDB species-representatives (release 207) after clustering at 99% sequence identity, to remove redundancy. NCBI-style taxdump files (nodes.dmp, names.dmp and accession2taxid) were then generated using scripts from https://github.com/nick-youngblut/.

Open reading frames were predicted on raw HiFi reads using FragGeneScan v1.31. Gene sequences were aligned to the generated GTDB protein database using Diamond blastp (parameters: --id 50 --top 5 --fast). A secondary filtering step was applied to the output including identity threshold >65%, e-value <1E-10 and query-cover >50%. Using the remaining hits, a single taxonomic classification for each gene was determined using a last common ancestor approach, *lca* command from TaxonKit [30]. To further increase the number of genes taxonomically classified, the last common ancestor algorithm was then applied to all genes within a single HiFi read, resulting in a single taxonomic classification for each HiFi read, and it’s containing genes.

### HiFi read functional annotation

For functional characterization, the predicted gene sequences (see above) from HiFi reads were subject to a custom annotation pipeline, modified from Priest *et al.* [31]. In brief, genes were annotated against the Pfam database (release 35.0) using HMMsearch VXXX (parameters: cut_ga), UniProtKB database (05.2022) using Diamond blastp (v2; parameters: -k 1 --evalue 1e- 10 --query-cover 50 --id 40 --sensitive) and the KEGG database (07.2022) using kofam_scan (https://github.com/takaram/kofam_scan ; parameters: -E 0.0001). Additional annotations were obtained by searching more specialised databases using HMMScan (Transporter Classification database; obtained 11.2021, TonB HMM profiles from TIGRFAM, and dbCAN; v10) and Diamond blastp (CAZyDB; release 09242021, SulfAtlas; v1.3, MEROPS; v12.1) using the same settings as described above, except for HMMScan against the dbCAN database (parameters; -E 1E-15).

### Single-copy ribosomal protein (SC-RBP) gene analysis

From the gene annotations, 16 single-copy ribosomal protein (SC-RBP) genes [32] were identified and extracted from each metagenome. The average number of SC-RBP was used as a proxy for the number of genomes recovered in each metagenome. A subset of four SC-RBP genes (RBP L3, L4, L6 and S8) were clustered at previously defined gene-specific ANI thresholds [33] and the average number of clusters across the four was used as a proxy for the number of species captured. The composition of metagenomes and metatranscriptomes was compared based on the species clusters from the RBP L6 gene, selected due to its high recoverability and species delineation accuracy [33]. The taxonomy of each cluster was determined based on a majority vote between the taxonomy of the contained genes, derived from original HiFi read classifications.

### Assembly, binning and metagenome-assembled genome recovery

The assembly of Hifi reads was performed using MetaFlye v2.8 [34] (parameters: --meta –pacbio-hifi –hifi-error 0.01 –keep-haplotypes). Coverage information was obtain through mapping HiFi reads to assembled contigs using Minimap2 v2.1 (parameters: -x map-hifi –MD). Contigs were binned using Metabat2 [35]. The resulting bins were subject to manual refinement using the Anvi’o v7 [36] interactive interface to generate metagenome-assembled genomes (MAGs). MAGs were dereplicated at a 99% ANI threshold using dRep v3.2.2 (parameters: --comp 50 --con 5 --sa 0.99 --nc 0.6). The completeness and contamination of representative MAGs was estimated using CheckM v1.1.2 [37]. A two-pronged approached was used for taxonomic classification of MAGs, the classify_wf pipeline of GTDB-tk v1.0.2 [38] (Release 207) and the extraction of 16S rRNA gene sequences using Barrnap v0.9 [39] and classification against the SILVA_SSU_Ref138.1_NR99 database, following the same process described in ‘*Phylogenetic characterisation of communities’*. Of the species-representative MAGs, 84% contained a complete 16S rRNA gene and thus received dual taxonomies.

### MAG relative abundance estimation

The relative abundance of representative MAGs was determined using a similar approach to Orellana et al. [40]. In brief, reads were competitively recruited from each metagenome to the MAG representatives. Mapped reads were converted into depth values using Genomecov (-bga option) from the Bedtools package [41] and the 80% central truncated average of the sequencing depth (TAD) was determined using the ‘BedGraph.tad.rb’ script (option range 80) from the enveomics collection [42]. The relative abundance was then determined as the quotient between the TAD value and the number of microbial genomes captured in each metagenome, determined from SC-RBP genes.

### MAG functional characterisation

The functional characterisation of MAGs was performed following the same procedure described for the HiFi reads above except for an additional process of polysaccharide utilisation loci (PULs) detection. PULs were defined as genetic loci containing a SusC/SusD gene pair with two or more degradative CAZymes or the presence of at least three degradative CAZymes in close proximity (maximum six genes apart). PULs were manually inspected and visualised at <u>BioRender.com</u>.

### Transcription level of genes at the community- and MAG-level

Adapters and low quality reads were removed from the metatranscriptomes using BBDuk of the BBtools program v38.73 (http://bbtools.jgi.doe.gov) (parameters: ktrim=r, k=29, mink=12, hdist=1, tbo=t, tpe=t, qtrim=rl, trimq=20, minlength=100). Although an rRNA depletion step was performed prior to sequencing, it is expected that 5 – 15% of reads would still be related to rRNA. As such, SortMeRNA v2.0 [44] was used to filter out rRNA sequences from the dataset, with the SILVA SSU Ref 138 NR99 database as a reference. The transcription level of genes was determined by read recruitment of transcripts to the predicted gene sequences from the HiFi reads using BBmap (v35; parameters: minid=98 idfilter=98). Mapped read values were converted to transcripts per million (TPM), according to Wagner *et al.* [45]. For MAGs, transcripts were competitively recruited to all MAG genes, with same procedure as above, and the values were converted to TPM using the total number of transcripts recruited to the whole metagenome-predicted genes as the total transcript values. In order to compare the transcription level of MAGs across samples, we determined the average TPM value of the 16 SC-RBPs for each MAG in each sample, and took the quotient of this and the average TPM value of the 16 SC-RBPs in the whole sample – providing proportional transcription of all genomes recovered. To place MAG CAZyme gene family transcription into the context of the whole community, we performed an additional read recruitment step. First, we concatenated genes from all MAGs into files based on CAZyme gene family or sub-family annotations. Then, we identified all transcripts that mapped to the metagenomic read-predicted genes for each of these families and subsequently recruited them to the concatenated MAG gene files, using the same parameters assigned above. Based on the number of transcripts mapped, the relative proportion of CAZyme gene family transcription was determined for each MAG.

## RESULTS & DISCUSSION

Seawater samples were collected from surface waters (SRF) and the bottom of the surface mixed layer (BML) in the Eastern Fram Strait region to investigate the distribution of carbohydrates and their utilisation by microbial communities. The ten sites were grouped into three categories based on the underlying seafloor topography (above-slope, above-shelf and open-ocean), which also correspond to differences in hydrographic conditions. The main water mass in this region is of North Atlantic origin. The West Spitsbergen Current (WSC) transports Atlantic water northward into the Arctic Ocean, with the main branch flowing above the continental slope. At the shelf break, a temperature-salinity front occurs, whilst above the West Spitsbergen shelf, Atlantic water (AW) converges and mixes with Arctic water and freshwater from land, resulting in intra-annual variability in hydrographic properties [46]. Based on temperature and salinity values, the main water masses in this region can be distinguished, with AW characterised by >34.9 psu and >4.1 °C [47]. The temperature of SRF and BML depths during sampling in this study were indicative of AW, ranging from 4.1 – 7.7 °C. However, the salinity values in SRF waters of above-shelf (S10) and three above-slope stations (S1, S6 and S8) were below the AW-defining thresholds (Figure 1 and Supplementary Table S1). These observations suggest an influence of either polar-derived water or freshwater from Spitsbergen at these stations.

### Carbohydrate analysis of POM samples

The monosaccharide and glycan composition of carbohydrates in POM (>0.7 µm) was analysed in SRF and BML depths at nine stations. The monosaccharide composition of carbohydrates in POM showed depth- and location-related patterns (Figure 2). Total neutral and amino monosaccharide concentrations ranged from 1.4 – 13.8 µg per L of seawater (hereon µg l^-1^). Higher values were typically observed in SRF, average of 8.8 µg l^-1^, compared to BML depths, average of 5.2 µg l^-1^ (Supplementary Table S3). However, the magnitude of change between the two depths was station-dependent, from a negligible difference at station S8 to a threefold decrease from SRF to BML depths at station S6 (Figure 2a). The decrease in monosaccharide concentrations with depth is in agreement with previous observations from the Pacific Ocean [48].

**Figure 2.**
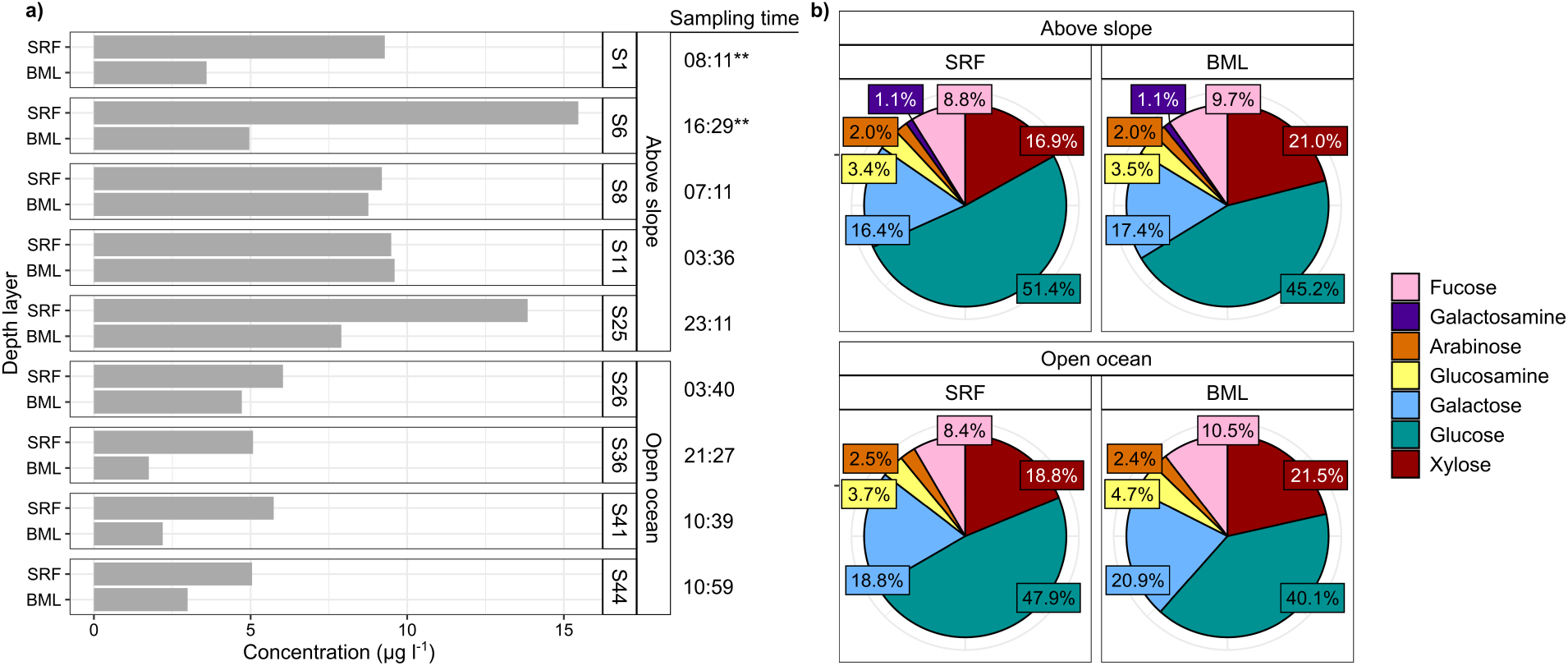
Total concentration and relative abundance of monosaccharides in carbohydrates from POM fraction. **a)** Total monosaccharide concentrations as ug per L of seawater at each station and depth**, b)** Relative abundance of monosaccharides grouped by station location in relation to continental slope. Data show neutral and amino monosaccharides. SRF = Surface water sample, BML = sample from bottom of the surface mixed layer.

In addition, above-slope stations contained higher monosaccharide concentrations than open-ocean stations (Figure 2 and Supplementary Figure S1). This spatial heterogeneity resembles that of chlorophyll *a* and dissolved organic compounds during early summer in this region, which reach highest concentrations in SRF depths above the continental slope (∼8 °E) [49]. These patterns likely reflect hydrographic processes, such as the frontal zone situated above the shelf break.

The most abundant monosaccharide detected in all samples was glucose. Glucose represented a larger relative proportion of POM carbohydrates in SRF (∼49%) than BML (∼43%) depths and in above-slope (∼47%) compared to open-ocean (∼44%) samples (Figure 2b). These values are within the range of those previously reported from oceanic surface waters (31 – 55%) [50–52] but lower than during phytoplankton blooms, wherein glucose can constitute >70% of POM carbohydrates [53]. Furthermore, in contrast to findings from the high North Pacific [50], the relative proportion of glucose decreased with depth, concurrent with an increase in all other monosaccharides. In particular, xylose increased 5% in relative proportion to the other monomers from SRF to BML depths (Figure 2b). The relative decrease in glucose indicates selective utilisation of glucose-containing glycans in surface waters.

Combining carbohydrate microarrays with monoclonal antibodies and carbohydrate binding modules resulted in the detection of 17 distinct glycan epitopes in POM (Supplementary Figure S2 and Supplementary Table S4). This structure-based detection provides semi-quantitative presence and abundance of distinct epitopes, where changes in antibody binding signal correlates to epitope abundance [28, 54]. Variations in the abundance of glycan epitopes exposed location and sample-specific patterns. The glycan epitopes observed most frequently included fucose-containing sulfated polysaccharide (FCSP) in 94% and glucuronoxylan in 89% of samples (Figure 3). FCSPs are unique to brown algae, wherein they serve important structural roles [55] and formulate part of the secreted carbon pool [56], while glucuronoxylans are common features of land plant and macroalgal cell walls [57]. Recently, we discovered both of these complex polysaccharides in microalgal blooms [28] as well as the secretion of FCSPs by diatoms in culture [58]. FCSPs synthesized by diatoms accumulate in POM over a period of weeks during a spring phytoplankton bloom, indicating stability against bacterial degradation [28], and can contribute to long-term carbon sequestration in sediments [59]. This observed stability contrasts β-1,3-glucans, such as laminarin, that are also synthesised by brown algae and diatoms. Laminarin is structurally simpler than FCSPs, with fewer unique linkages and without sulfate esters. Concurrently, the presence and activity of laminarases is observed more frequently in marine microbes compared to those targeting FCSPs, suggesting higher consumption and turnover [10, 28, 60]. In our samples, laminarin was absent at some open-ocean stations, but present in all above-slope sites (Supplementary Figure S2). In addition to a potentially more rapidutilisation by microbes, the heterogeneous distribution of laminarin may also result from variations in the distribution of phytoplankton, reflecting previous observations from this region [61].

Sampling conducted at the same location over a two day period (stations S1 and S6) and into deeper waters (down to 200 m) showed additional vertical as well as temporal shifts in the abundance and diversity of POM carbohydrates (Supplementary Figure S4). S1 samples were retrieved at 08:00 whilst S6 samples were collected one day later at 16:30. The absolute concentration of monosaccharides in POM carbohydrates was 1.1 – 3.6x higher in S6 compared to S1 samples. Temporal shifts were also evident in POM glycans. Notably, SRF depths in S6 samples contained more α-1,5-arabinan and β-1,3-glucan as well as β-1,4-xylan epitopes and xylosyl residues. The only epitopes more abundant in S1 samples were alginate and glucuronoxylan. It is important to note that the alginate targeting antibody used, BAM7, also has cross reactivity with FCSP. The pronounced shift in glucose and β-1,3-glucan between the two sampling time points may reflect diurnal fluctuations in laminarin production by diatoms (Becker et al. 2020), with highest production during the day and a partial consumption at night. Such diel periodicity is also evident at the particulate organic carbon level [62], with accumulations during the day to a maximum concentration at dusk [63]. In addition, the pronounced increase in xylosyl residues and β-1,4-xylan epitopes at S6 could suggest the introduction of, or change in the primary producers between the two sampling time points, potentially resulting from shifts in water mass dynamics.

**Figure 3.**
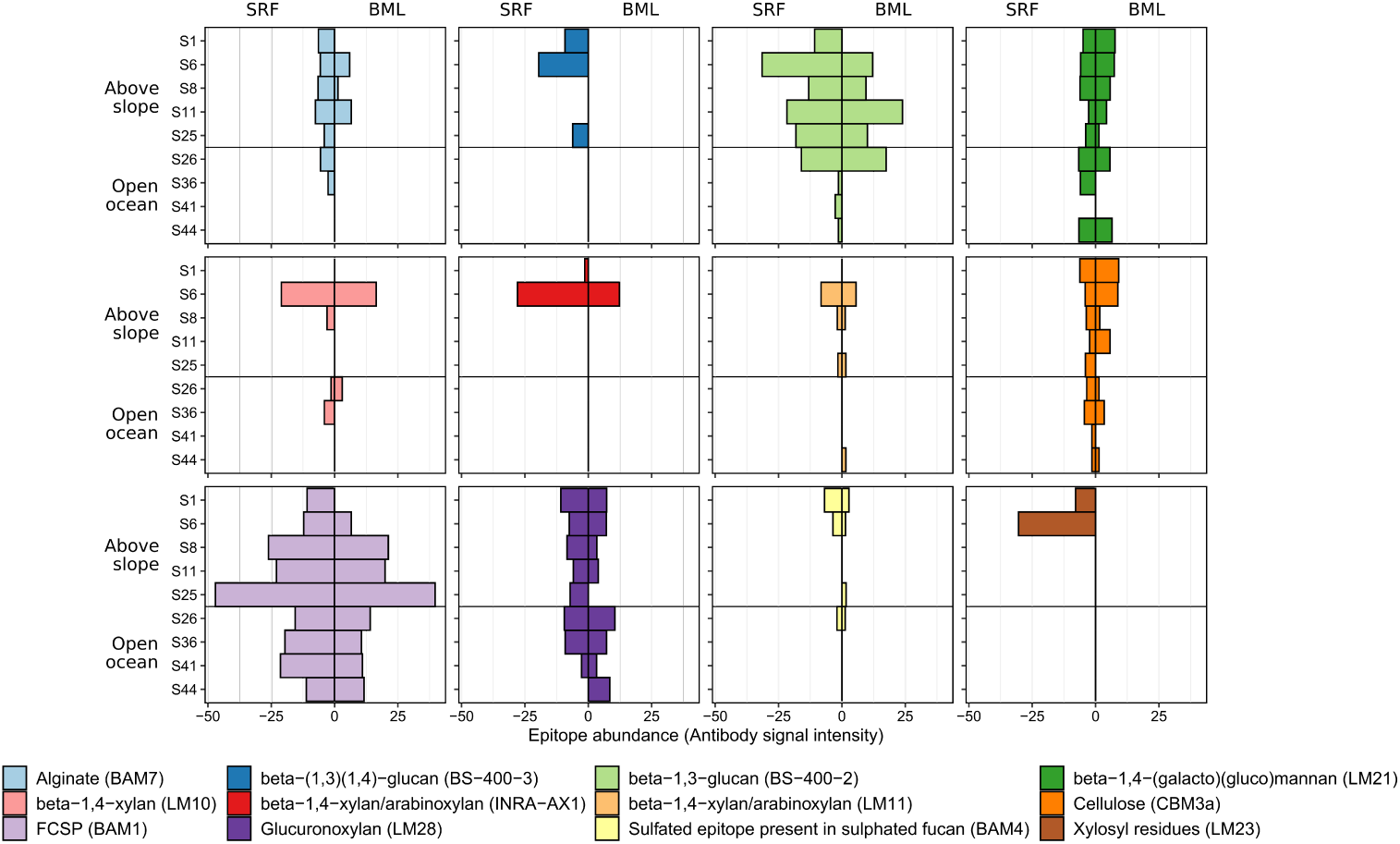
Diverse glycan structures occur in Atlantic waters of the Arctic. Plots show the relative abundance (antibody signal intensity) of glycan epitopes detected in POM. Signal intensities for each extract against each antibody were quantified and the highest mean signal value (n=4 replicates per extract) in the data set (which corresponded to a standard control) was set to 100 and all other values were normalised accordingly. The epitope abundances shown here represent the summed values from the mean signal of H_2_O, EDTA and NaOH extractions. Each antibody has a different avidity, thus the absolute numbers should not be compared between probes to infer different concentrations, but signal from a single antibody determines the relative abundance of the epitope within the sample set. SRF = Surface water sample, BML = bottom of the surface mixed layer. FCSP = Fucose-containing sulphated glycan. Some epitopes that were only detected at low intensity in <3 samples were not included in this plot.

### Sampled microbial communities were indicative of summer in high-latitude Atlantic waters

To assess microbial carbohydrate utilisation patterns, eight PacBio HiFi read metagenomes were generated from SRF and BML depths of two above-slope (S8 and S25), one open-ocean (S26) and one above-shelf (S10) station. From four of the samples, metatranscriptomes were also generated (Supplementary Table S5 and S6). Although samples for carbohydrate characterisation were not obtained for the above-shelf station S10, the lower salinity values observed in these samples (psu < 33) indicates influence from polar marine/freshwater and thus predictably provides a valuable comparison.

We performed a taxonomy-independent comparison of the sequenced communities here with previously published metagenomes from the Fram Strait [31] and Arctic Ocean [64]. Based on sequence composition dissimilarity, our metagenomes were most closely related to those previously generated from WSC, high North Atlantic and Barents Sea samples in June and July and most dissimilar to those from the polar water mass of the western Fram Strait (Supplementary Figure S5). This contrast indicates communities captured by our metagenomes, derived from September sampling, are representative of summer communities in Atlantic waters of the Arctic.

To focus explicitly on microbial carbohydrate utilisation, we removed metagenomic reads not classified as Bacteria or Archaea. Despite the size fractionation employed during sampling (0.2 – 3 µm), 22– 49% of the metagenomics reads were classified as Eukarya. Although our analysis was concentrated on the prokaryotic fraction, we also extracted and analysed 18S rRNA genes to provide insights into the eukaryotic taxa present in samples (Supplementary Figure S7). Using the average sequencing depth of single-copy ribosomal protein (SC-RBP) genes, we determined the number of microbial genomes sequenced to range from 761 in S8_SRF to 1619 in S10_BML (Supplementary Figure S6b). The number of genomes detected is used to normalize the abundance of functional genes.

### Composition of metagenome and metatranscriptome microbial communities

The composition and structure of microbial communities varied across samples, inconsistently with changes in location or depth. The most prominent families across the metagenomes were the *Flavobacteriaceae* (4 – 14%), *Rhodobacteraceae* (3 – 13%), D2472 (SAR86; 5 – 12%) and *Poseidoniaceae* (2 – 12%) except for in S25_BML, which was enriched in *Alteromonadaceae* (25%) (Supplementary Figure S8). These bacterial families were also substantial contributors to community transcription. However, the relative proportions of abundance and transcription were not consistent, e.g. *Flavobacteriaceae* represented a two- to threefold higher proportion of transcription. Twenty genera were identified as reaching >2.5% relative abundance and together, constituted ∼42% of the microbial communities (Figure 4). These 20 genera included *Pseudoalteromonas* (0 – 29%), MGIIa-L1 (<0.1 - 10%), *Amylibacter* (<0.1 – 6%), D2472 (SAR86; 3 – 6%), ASP10-02a (<0.1 – 6%) and HTCC2207 (SAR92; 1 – 6%). However, only some of these genera were substantial contributors to community transcription. The maximum relative proportion of transcription was observed in *Pseudoalteromonas* (45%), *Vibrio* (29%), *Flavobacterium* (13%), *Amylibacter* (9%), SAR86 (9%) and *Pseudothioglobus* (9%). Discrepancies in the community structures between metagenomes and metatranscriptomes highlight that gene abundance is not always intrinsically linked to transcription. This disconnect was particularly evident for MGIIa-L1 (Marine Group II Archaea), which represented a larger proportion of gene abundance than transcription, and genera within the *Bacteroidia*, such as *Flavobacterium* and MAG121220-bin8 (NS4), which showed the opposite trend. Such patterns are not uncommon in marine microbial communities [65] and can be influenced by lifestyle, fitness and differences in metabolism.

**Figure 4.**
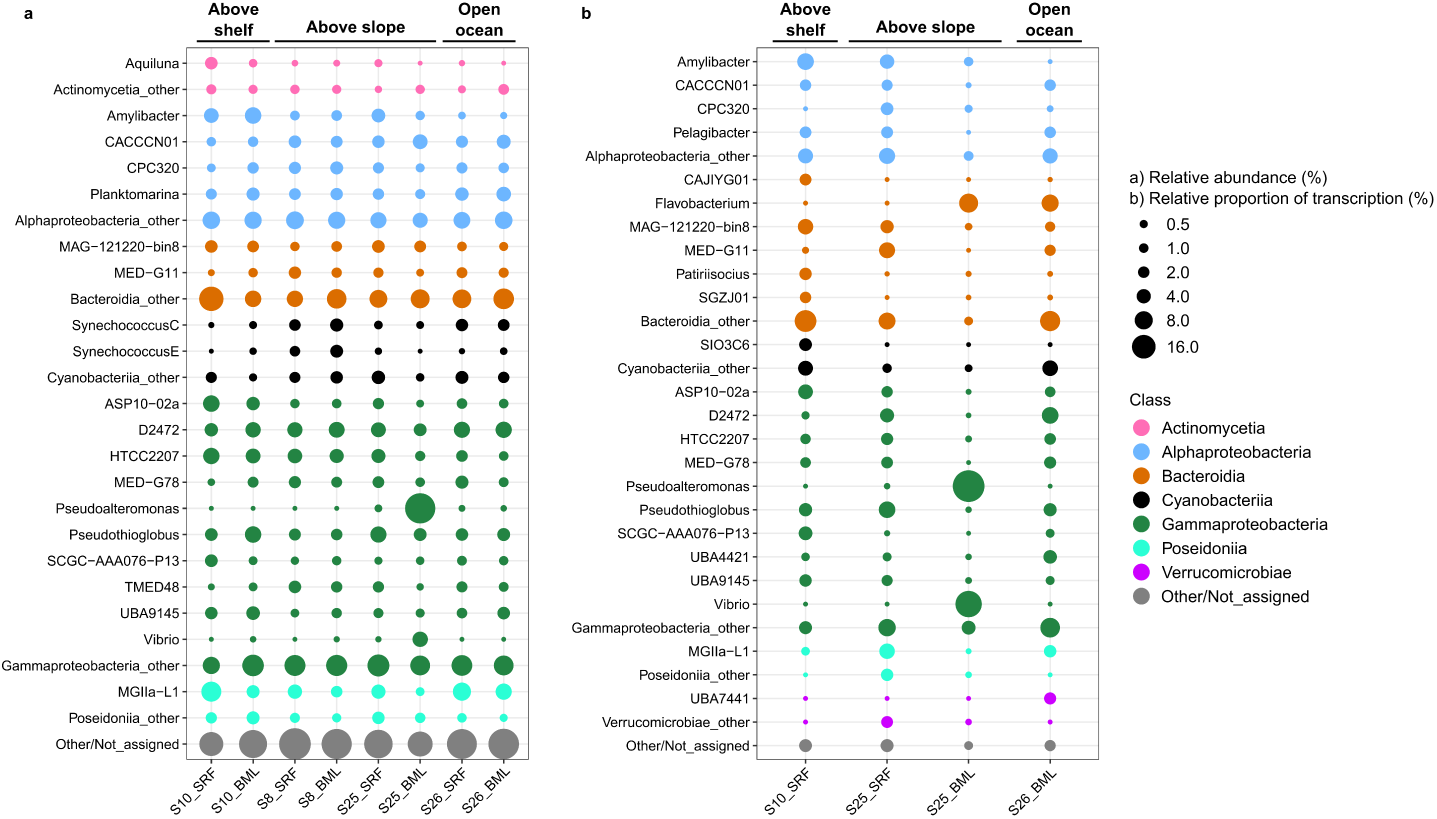
Composition of microbial communities in metagenomes and metatranscriptomes were sample-specific. Relative abundance is described as the relative proportion to total ribosomal protein L6 sequences identified in the sample. Relative proportion of transcription is determined as the relative proportion to total sample ribosomal protein L6 TPM values. Taxonomy of ribosomal protein L6 sequences was derived from taxonomic classification of the original HiFi read, which was classified using a GTDB-based database.

Sample S25_BML harboured a distinct community from all others, with a large proportion of *Pseudoaltermonas* and *Vibrio.* These genera have previously been observed under nutrient-rich conditions and are known to be associated with eukaryotic hosts and phytoplankton blooms [66, 67]. Considering the high proportion of eukaryotic reads captured in some of the metagenomes, the observed pattern in S25_BML may represent signals of processes occurring in the larger size fraction. In support of this, the eukaryotic community of this sample (Supplementary Figure S7) contained a large proportion of 18S rRNA genes affiliated with copepods, whose microbiomes are often enriched with *Pseudoalteromonas* and *Vibrio* [68, 69].

In general, the microbial community compositions observed here more closely resemble those previously described from summer [70, 71] than from late September in the same region [72]. This difference likely reflects the inter-annual variability in seasonal transitions, as September separates summer from the beginning of winter. However, methodological differences also play a major role. Methodological influence is particularly evident with respect to the high proportions of MGIIa-L1 in our samples, which has not been observed before in the WSC, and is typical of late summer communities in temperate coastal ecosystems [73]. The previous employment of bacterial-specific 16S rRNA gene based primers [71] has likely contributed to the MGII, and Archaea more generally, being overlooked in the WSC, wherein they could represent an important fraction of the microbial community.

### Carbohydrate utilisation patterns of microbial communities were sample-specific

Microbial carbohydrate utilisation was assessed through the abundance and transcription of CAZymes. In particular, we focused on those involved in degradative processes (glycoside hydrolase, GH; carbohydrate-binding module, CBM; carbohydrate esterase, CE; polysaccharide lyase, PL).

CAZyme genes represented a minor proportion of community gene content but a higher proportion of gene transcription. The number of CAZyme genes ranged from 7 – 19 per microbial genome (PMG), which corresponded to 0.3% of community gene content, on average. In contrast, CAZyme genes represented between 1.5% and 3.0% of community gene transcription. Clear differences were also observed with respect to the number of CAZyme genes being transcribed, with 3159 genes in S10_SRF compared to 494 genes in S25_BML. Employing a dissimilarity-based approach, we observed that the CAZyme compositions of samples from S8 and S26 were grouped together based on station (location) whereas those of S10 and S25 showed no coherent clustering (Supplementary Figure S9).

The CAZyme gene profiles comprised a core backbone of universally abundant and transcribed gene families. The most abundant gene families in all samples were those involved in peptidoglycan synthesis and degradation, which is in agreement with previous observations and reflects the core machinery required for bacterial cell membrane construction and maintenance [21]. Core CAZyme gene families included CE11, GH23, GH103 and GH73 that together represented, on average, 27% of CAZyme gene abundance (average of 0.83 PMG) and 35% of CAZyme gene transcription (Figure 5 and Supplementary Tables S7 – S10). Several CAZyme gene families involved in the degradation of algal-derived glycans also represented a high proportion of gene abundance and transcription. The GH16_3 gene family, which contains enzymes that degrade laminarin [74], constituted 3.1% of CAZyme gene abundance (0.3 PMG) and 2.7% of CAZyme gene transcription. Other prominent gene families included those known to target sialic acids (GH33 [75]; ∼1.4% of CAZyme abundance and 1.7% of CAZyme transcription), α-mannans (GH92 [76]; ∼1.7% CAZyme abundance and ∼1.3% CAZyme transcription) and alpha-linked fucose that is common in FCSPs [77] (GH29; ∼1.3% CAZyme abundance and 1.2% CAZyme transcription) (Figure 5 and Supplementary Tables S7 – S10). Positive linear relationships between CAZyme gene family abundance and transcription was observed in all samples except S25_BML (Supplementary Figure S10) – a higher abundance resulted in higher transcription. However, exceptions to this correlation were observed, with some CAZyme families exhibiting nearly twofold higher relative transcription than abundance, such as the alginate-targeting PL7_5 [78] in S10_SRF and the galacturonan-targeting GH28 [79] in S26_BML (Supplementary Figure S11).

**Figure 5.**
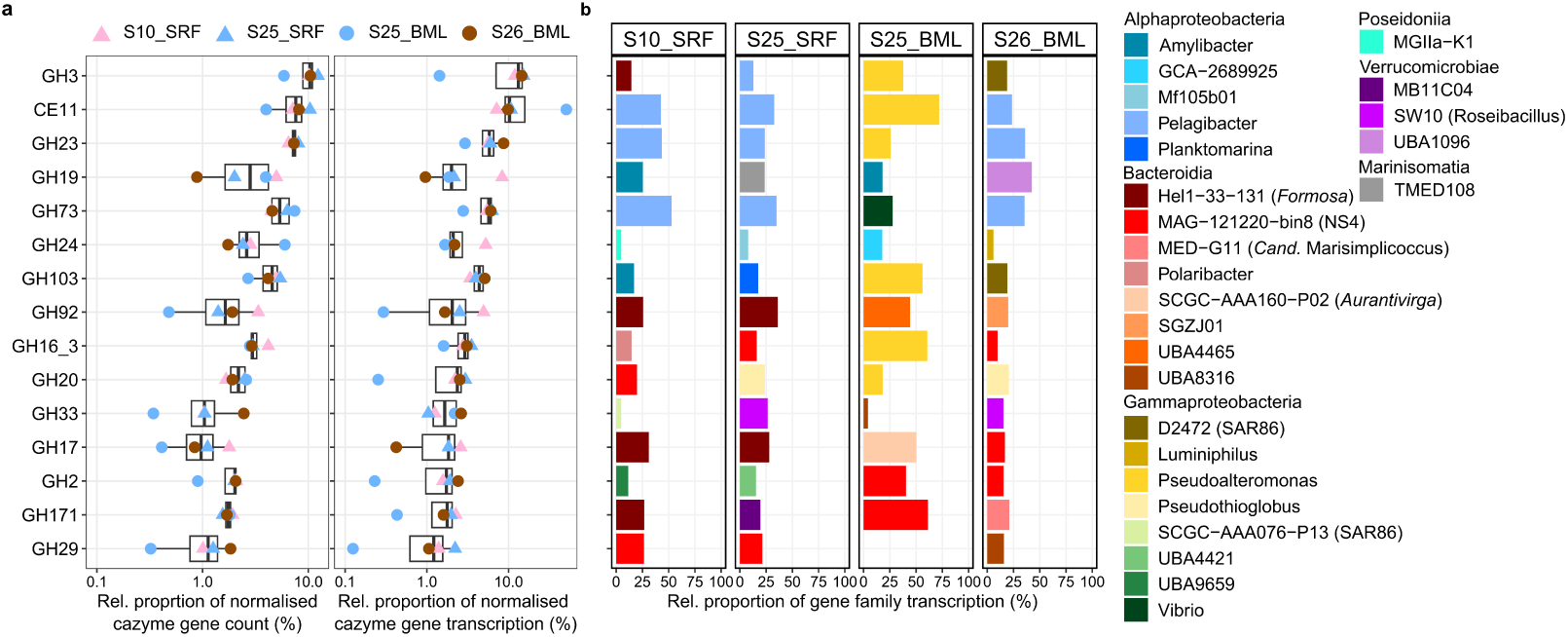
Abundance, transcription and taxonomic information of the CAZyme gene families with the highest proportional transcription across samples. **a)** The abundance and transcription of carbohydrate-active enzyme (CAZyme) genes was normalised by the number of microbial genomes per sample and then converted to relative proportions. The selected CAZyme gene families visualised are those that reached the highest proportions of transcription across samples. **b)** The genera that contributed the most to transcription of each gene family in each metatranscriptome. Taxonomic classification of CAZyme genes was derived from the classification of the read from which it was derived.

Although the microbial community analysis was concentrated on the free-living fraction, corresponding patterns were observed between CAZyme gene transcription and their target glycans in POM. For example, the widespread presence of FCSPs and alginate and the transcription of alpha-fucosidases (GH29) and alginate lyases (PL7_5). Glycans that are part of the algae cell belong to the POM pool, but can become part of DOM through cell lysis, viral infection and grazing (a likely process given the presence of copepods in the extracted 18S rRNA gene data). The same glycan epitopes can thus be present in both POM and DOM, as has been evidenced during phytoplankton blooms [28]. Therefore, the glycans detected in POM here, were likely also available to free-living heterotrophic microbes.

Distinct microbial taxa were responsible for the transcription of CAZyme families in each sample. For CAZyme gene families involved in bacterial glycan recycling, which are universally present in microbes, the transcription was dominated by the most abundant taxa in the community, with *Pelagibacter,* accounting for ∼45% of the peptidoglycan-targeting GH23 family transcription (Figure 5). In contrast, the taxa contributing the most to transcription of algal glycan-targeting CAZyme families were affiliated with *Bacteroidia*, including *Polaribacter* and NS4 for laminarin (GH16_3), *Formosa* for α-mannan (GH92) and NS4 and UBA8316 for FCSP (GH29). Several of these genera are well known as carbohydrate-degrading specialists in temperate coastal ecosystems and are among the main microbial responders to spring phytoplankton blooms [21, 80]. In particular, *Polaribacter* and *Formosa* have been identified as annually recurring after phytoplankton blooms in the German Bight, North Sea, reaching > 5% relative abundance, and exhibiting successional-like dynamics indicative of glycan-based niche partitioning [10, 21]. The presence and activity of these genera in our samples indicates that they are also key players in carbohydrate cycling in high-latitude waters and at later seasonal stages. To investigate carbohydrate utilisation by microbes at a higher resolution and assess whether glycan-based niche partitioning occurs, we analysed patterns at the population-level using metagenome-assembled genomes.

### Recovery of metagenome-assembled genomes

We recovered and analysed 83 population-representative metagenome-assembled genomes (MAGs), delineated at a 99% ANI threshold (Supplementary Information S1 and Supplementary Table S8). The MAGs captured between 48 – 88% of the microbial metagenomic reads (Supplementary Table S12) and 11 – 37% of the microbial metatranscriptomic reads (Supplementary Table S13) (Figure 6). Analogous to the community-level patterns, the coupling of relative abundance and relative proportion of transcription varied across MAGs and taxa. MAGs affiliated with MGIIa-L1 (Marine Group II Archaea) exhibited high relative abundance (up to 12%) but low relative transcription (up to 1.5%) while the *Aurantivirga*-affiliated MAG showed the opposite trend, with ∼0.7% relative abundance but ∼2.7% relative proportion of transcription. A MAG (S25_BML_bin_129) that shares 97.1% ANI to *Pseudoalteromonas primoryensis* was also recovered. The *Pseudoalteromonas primoryensis* MAG constituted 22.8% relative abundance of sample S25_BML, indicating that the pattern observed at the community-level was driven by a single species.

**Figure 6.**
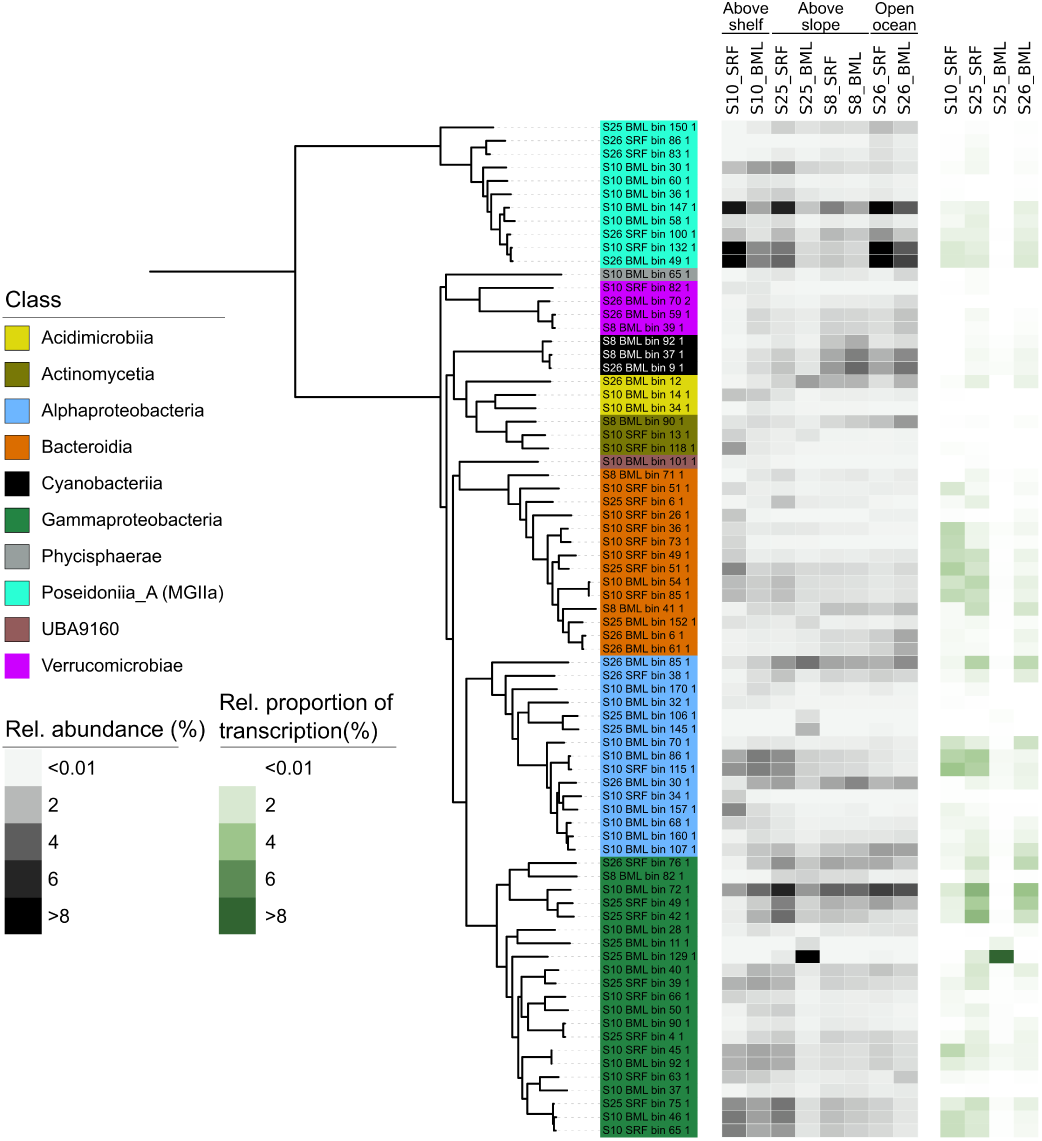
Microbial populations show distinct patterns in distribution and activity across samples. The tree was calculated from a concatenated alignment of 16 single-copy ribosomal protein genes (SC-RBPs), with a threshold of at least 8 genes per MAG for inclusion in the tree. Relative abundance of MAGs was defined as the quotient between the truncated average sequencing depth of the MAG and the average sequencing depth of 16 SC-RBPs in the sample. Relative proportion of transcription was defined by the average TPM value of the MAG-encoded 16 SC-RBPs and the whole sample average TPM values for the 16 SC-RBPs.

### Sample-specific carbohydrate utilisation by individual microbial populations

Carbohydrate utilisation patterns in populations was assessed through the abundance and transcription of CAZymes, TonB-Dependent Transporters (TBDTs), sulfatases and peptidases. Peptidases were included for comparison as proteins are another key substrate used by heterotrophic microbes. The largest CAZyme gene repertoires were observed in *Verrucomicrobiae*- and *Bacteroidia*-affiliated MAGs, with an average of 15 and 14 CAZymes per Mbp, respectively (Figure 7). In contrast, *Poseidoniia*-affiliated MAGs harboured few CAZymes, 1 per Mbp, but high peptidase:CAZyme ratios, ∼3.7:1. The high peptidase content of *Poseidoniia*- MAGs indicates a preference for proteinaceous substrates, inline with previous observations for this taxa [73]. With respect to sulfatases, *Verrucomicrobiae*-affiliated MAGs harboured the most extensive repertoires, with an average of 24 per Mbp. Despite these taxa-related patterns, large variations in CAZyme repertoires were observed across MAGs, with a range of 5 to 25 per Mbp observed in *Bacteroidia* (Figure 7). The observed differences in carbohydrate degradation potential for these taxa are in accordance with previous findings, with large CAZyme repertoires reported for *Bacteroidia* and a specialisation on sulfated polysaccharides in *Verrucomicrobiae* [40, 77].

**Figure 7.**
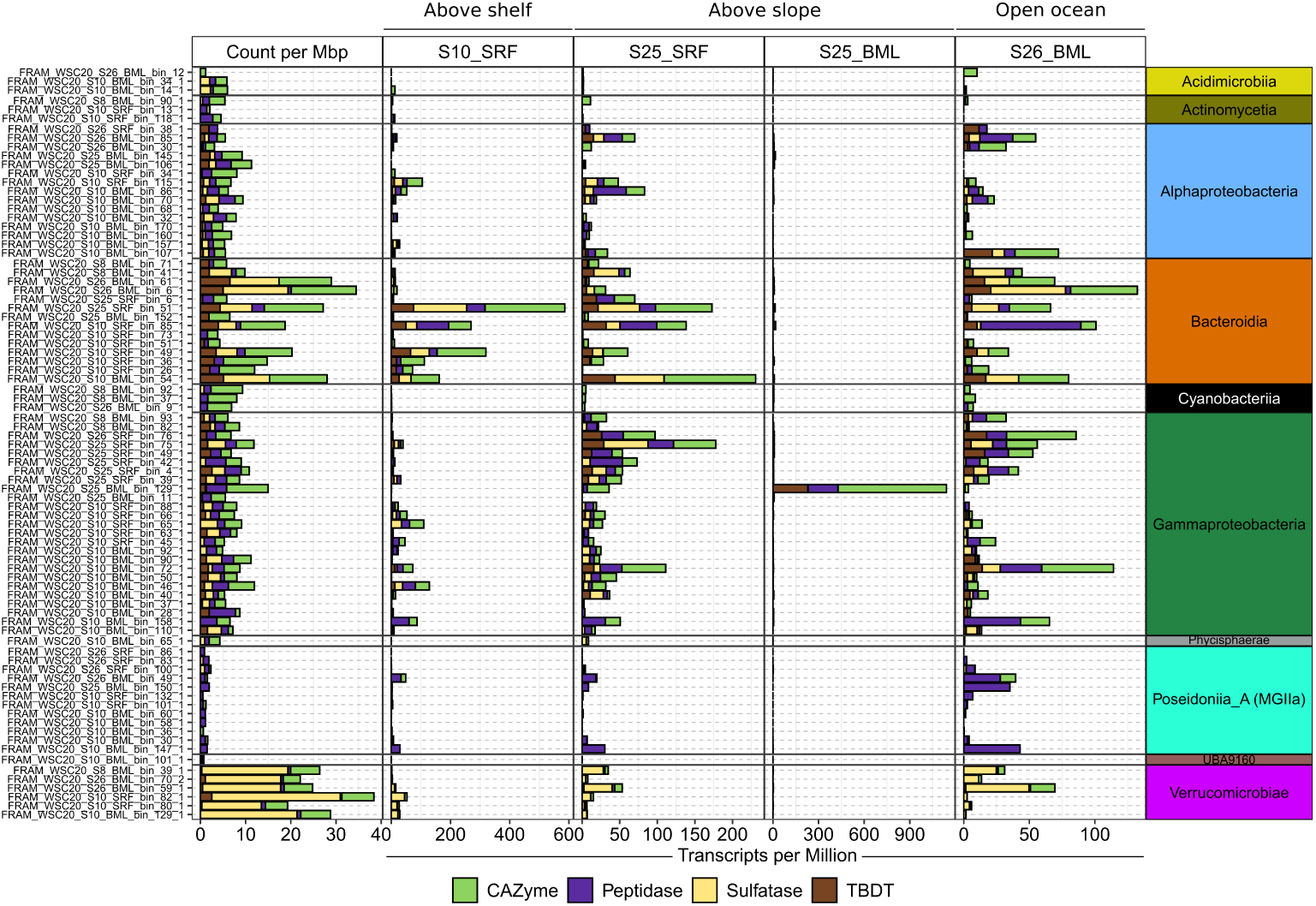
Count and transcription level of carbohydrate utilisation genes for population-representative MAGs. Carbohydrate utilisation genes were clustered into four groups based on function. The count and TPM value of all genes within each group were summed. Gene counts were further normalised by MAG genome size. TBDT = TonB-dependent transporter.

Comparing transcription levels of the focal gene groups across MAGs revealed distinct assemblages of populations that dominated carbohydrate utilisation in each sample (Figure 7). In the above-shelf sample S10_SRF, CAZyme, sulfatase and TBDT transcription values were dominated by only a few *Bacteroidia* representatives, particularly *Formosa* (25_SRF_bin_51_1) and NS2b (S10_SRF_bin_49_1). In contrast, the above-slope sample S25_SRF was characterised by a larger number and diversity of populations of the *Bacteroidia* and *Gammaproteobacteria* that exhibited comparable transcription values. The key contributors to CAZyme transcription in S25_SRF included two distinct populations affiliated with the NS4 Marine Group (S10_SRF_bin_85_1 and S10_BML_bin_54_1) along with one of *Formosa* (S25_SRF_bin_51_1), SAR92 (S25_SRF_bin_75_1) and SAR86 (S10_BML_bin_72_1). In addition, high sulfatase transcription in S25_SRF was observed for two populations of *Roseibacillus* (*Verrucomicrobiae*). In sample S26_BML, comparably high CAZyme gene transcription levels were observed across numerous populations assigned to *Bacteroidia, Gammaproteobacteria, Poseidoniia, Alphaproteobacteria* and *Verrucomicrobiae* that were less active in the other samples, which may reflect differences in depth. The largest contributors to CAZyme transcription in S26_BML that were less active in other samples included those assigned to *Cand.* Arcticimaribacter (S26_BML_bin_6_1) [81], *Flavobacteriaceae* (S26_BML_bin_6_1) and *Planktomarina* (S10_BML_bin_107_1). As could be expected, the TPM values for the focal gene groups in S25_BML were strongly dominated by the *Pseudoalteromonas* representative. The MAG-based analysis revealed that carbohydrate utilisation by populations is highly dynamic across samples, and thus heterogeneous over spatial scales. Furthermore, the comparably higher transcription of carbohydrate-degradation related genes by certain MAGs in each sample, suggests that the community-level patterns observed may be driven by only a minority of populations.

### Microbial populations make use of communal and specialist glycans

To place population transcription into the context of the communities, we determined the proportion of community transcription of each CAZyme gene family by each MAG (see Methods). We further focused on the top six MAGs contributing to transcription of each gene family and within those MAGs, the gene families that were more transcribed than single-copy ribosomal protein genes – considered up-transcribed in relation to the genome (Figure 8).

**Figure 8.**
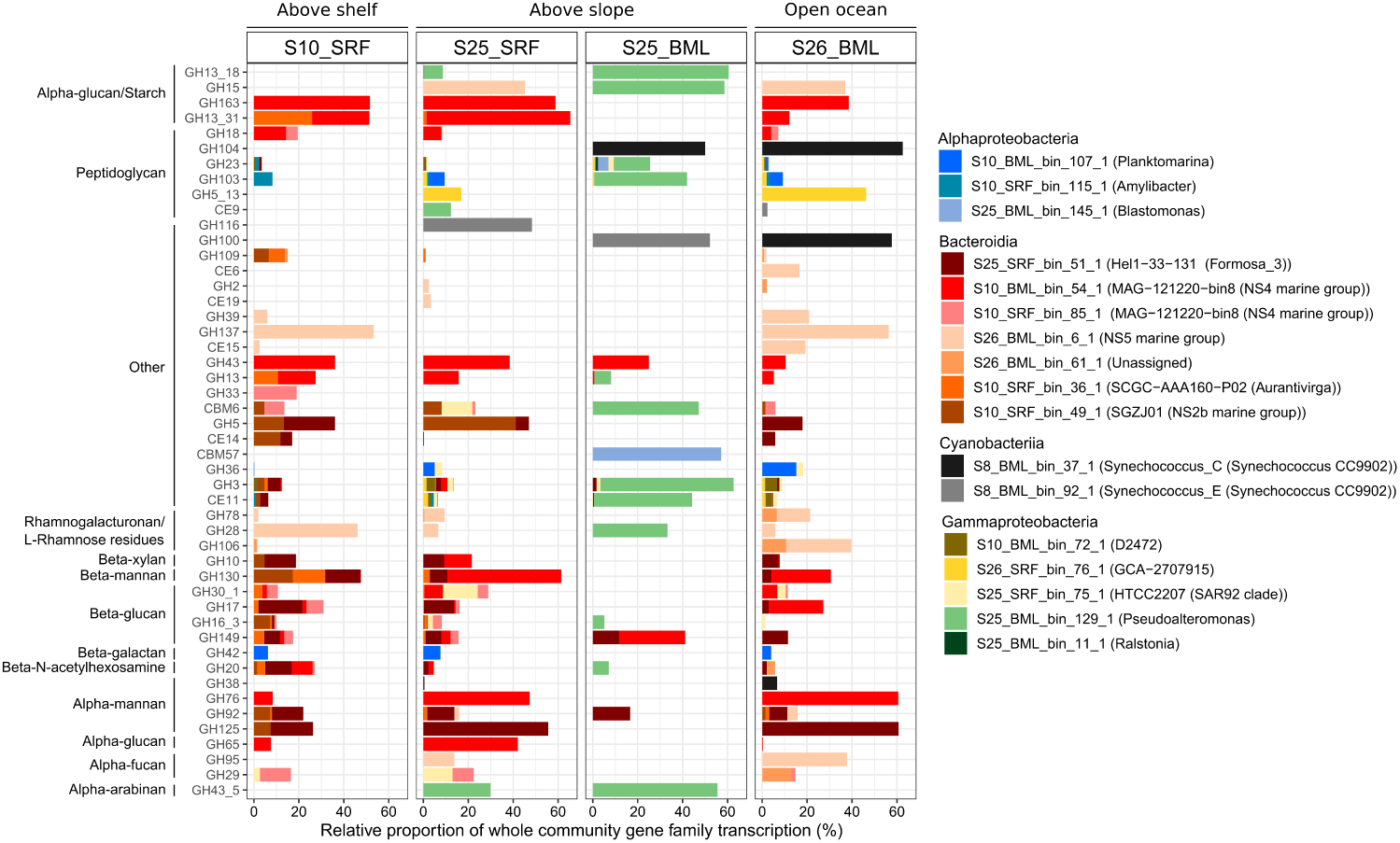
Comparison of CAZyme gene family transcription by selected MAGs across the four metatranscriptome samples. The ten MAGs that exhibited the highest CAZyme transcription values in each sample were selected. Within each of those MAGs, only CAZyme gene families with transcription values higher than ribosomal proteins were retained. To determine the proportion of transcription at the gene family level, we mapped all transcripts identified as CAZymes at the community level to MAG predicted genes. Presented here is the proportion of transcripts of each gene family associated with the selected MAGs – proportion of sample total transcription for that gene family.

Amongst the populations contributing the most to CAZyme gene transcription, we observed “communal” and “specialist” glycan utilisation patterns as well as glycan-based niche partitioning. The use of “communal” glycans was evidenced by several populations exhibiting similar proportions of transcription for a single CAZyme gene family in each sample and combined, only representing a fraction of total community transcription (Figure 8). The most notable “communal” glycans was laminarin (GH16_3 and GH17), which reflects how its simpler structure provides relatively high carbon energy with limited enzyme investment for microbes. In contrast, “specialist” glycans were evidenced by the transcription of a CAZyme gene being observed in only few populations and typically dominated by a single population in each sample. “Specialist” glycans included FCSP (GH29 and GH95) and α-mannan (GH76 [12] and GH125 [82]) and β-xylan (GH10 [83]). Populations targeting “specialist” glycans were typically affiliated with *Bacteroidia,* which also exhibited patterns of glycan-based niche partitioning, as has been observed before during phytoplankton blooms [22]. Glycan niche partitioning was observed when one population dominated the whole community transcription for a CAZyme gene family across multiple samples, such as rhamnogalacturonan (GH28 and GH78) by an NS5 population, or when a single, but distinct population dominated in each sample, as was observed for β-xylan (GH10) and β-mannan (GH130) (Figure 8). In addition, there were also cases where “specialist” glycans were targeted by two discrete *Bacteroidia* populations through different enzymes, including GH76 by an NS4 population and GH125 by a Formosa population that both target α-mannan. From this analysis, we evidenced the use of “communal” glycan substrates by multiple populations along with “specialist” glycans being targeted by either a single or a small number of unique populations across samples.

### Presence and transcription of polysaccharide utilisation loci

We further investigated carbohydrate utilisation by *Bacteroidia* populations through the transcription of their polysaccharide utilisation loci (PULs). PULs are the result of genomic rearrangement through evolution to optimise the regulation of genes involved in the degradation of a single glycan. Many specialist carbohydrate degraders of *Bacteroidia* harbour PULs. Investigating PUL compositions has been a valuable approach to disentangle glycan-based niche partitioning [10, 17, 22]. However, we identified PULs in only seven of the *Bacteroidia* MAGs recovered here (Supplementary Table S14 and S15), an observation that could be influenced by the completeness of the MAGs. Assessing the transcription of these PULs provided little additional insights into glycan utilisation patterns. In population MAGs with multiple PULs that target different substrates, transcription levels were comparable for each, indicating little regulation. Only in the case of the NS2b representative were distinct PUL transcription levels observed, with a xylan-targeting PUL exhibiting twofold higher transcription than a mannan-targeting PUL in S26_BML (Supplementary Table S14 and S15). This is an important observation for future studies investigating carbohydrate utilisation by members of *Bacteroidia*, as it indicates that the entire genomic repertoire of CAZymes, sulfatases and TBDTs needs to be considered and not only the information encapsulated within PULs. Furthermore, the lack of transcriptional variability across PULs indicates a low level of regulation and is more indicative of a priming effect, where the detection of one substrate results in transcription of all PULs.

## CONCLUSION

In Atlantic waters of the Arctic during late summer, the distribution of POM carbohydrates and their utilisation by microbes exhibit heterogeneous patterns over spatial scales. The monomeric and glycan composition of POM carbohydrates varied across locations and depths. Typically, higher abundances were observed above the continental slope compared to open-ocean locations. Monosaccharide compositions were dominated by glucose, which decreased in proportion with depth, suggesting preferential utilisation of glucose-based glycans in surface waters. Structurally complex glycans, such as FCSPs that accumulate in POM during phytoplankton blooms, were widely detected, while those with more simple structures, such as laminarin, exhibited patchy distributions. The observed distributions of POM carbohydrates is likely a result of spatial heterogeneity in primary production, as has been described from early summer in this region [49], along with variations in microbial utilisation. Through metatranscriptome analysis, we identified the active fraction of microbial communities and showed that unique assemblages of populations dominated carbohydrate utilisation in each sample. Prominent carbohydrate-degrading populations were found to make use of “communal” and “specialist” glycans. Those targeting “specialist” glycans, including *Formosa, Aurantivirga* and NS4 Marine Group populations, each exhibited distinct CAZyme transcription profiles across samples, reflecting glycan-based niche partitioning. These specialists taxonomically resembled, at the genus-level, those known to be primary responders and key carbohydrate degraders during phytoplankton blooms in temperate ecosystems. In combination, these results highlight that local biological and physical processes shape the POM carbohydrate pool in Atlantic waters of the Arctic during late summer, and that the key heterotrophic microbial populations actively respond to differences in glycan availability.

## DATA AVAILABILITY

The measurements of several abiotic parameters from sensors mounted on the CTD have been published under the PANGAEA accession 943220 [84]. The monosaccharide concentrations have been deposited under the PANGAEA accession 957737. The metagenomic raw reads, assemblies and metagenome-assembled genomes along with the metatranscriptomic raw reads were deposited at ENI-EBA, under the project accession PRJEB58071 (Supplementary Table S16). Detailed information on how to reproduce our results and generate the figures presented in this manuscript is provided at https://github.com/tpriest0/FRAM_STRAIT_WSC20_data_analysis.

## AUTHOR CONTRIBUTIONS

TP wrote the manuscript, performed the metagenomic, metatranscriptomic and carbohydrate data analysis. TP and SVM extracted the carbohydrates and SVM subsequently carried out the microarray analysis. SVM, JHH, BMF and RA contributed to the interpretation of the results and the formulation of the story. BMF, RA and TP planned the work and devised the project. All authors contributed to reviewing and improving the manuscript.

## ACKNOWLEDGEMENTS

We would like to thank the captain and crew of the *RV. Maria. S. Merian* for their support throughout all sampling aspects of this project. We thank Alek Bolte for his assistance with HPAEC-PAD. We thank Bruno Hüttel and the team at the Max Planck Genome Centre in Cologne for their work in generating the metagenomes and metatranscriptomes. We thank A. Murat Eren for his valuable comments and feedback with respect to data analysis. We thank the Max Planck Society for funding, and Hehemann further acknowledges funding from the DFG Heisenberg program HE 7217/5-1 and the DFG Exzellenzcluster 2077.

