## Supplementary Figures for "Spatial heterogeneity in carbohydrates and their utilisation by microbes in the high North Atlantic"

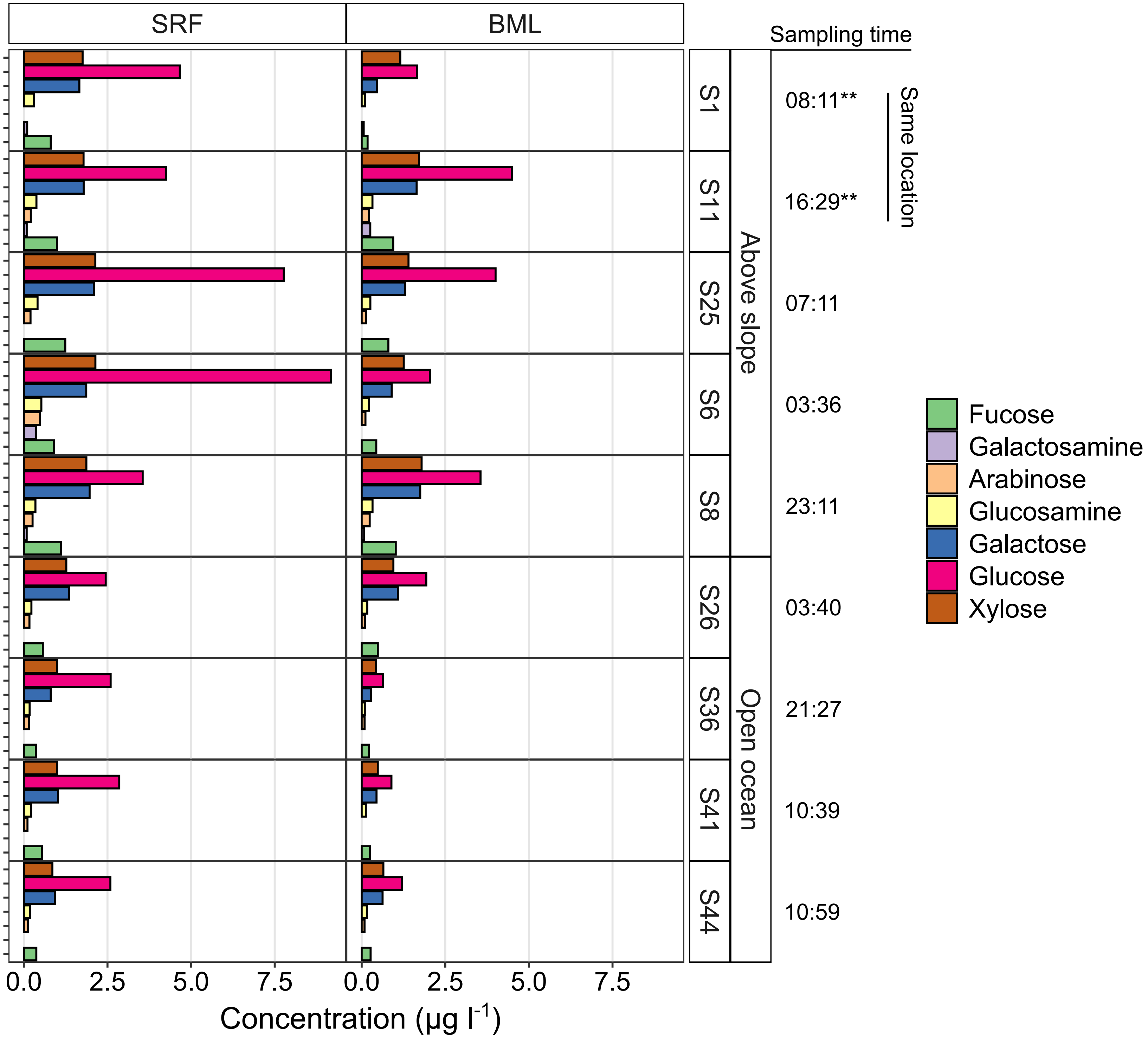


**Supplementary Figure S1. Monosaccharide concentrations in the particulate organic matter fraction.** Samples collected from the same location are indicated with ***. SRF = surface water, BML = bottom of surface mixed layer. Concentrations shown represent 'per L of sampled seawater'.

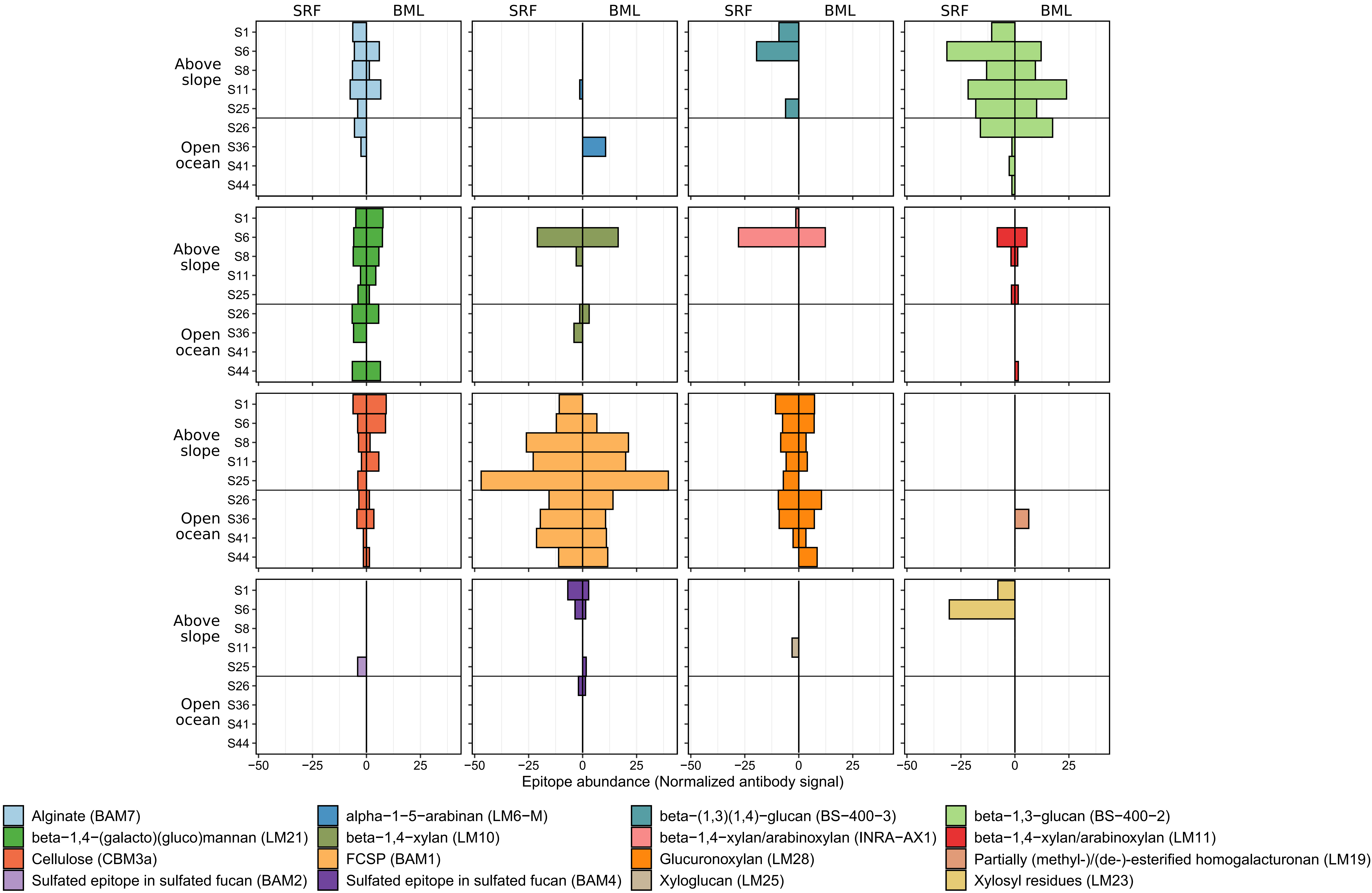


**Supplementary Figure S2. Relative abundance of polysaccharide epitopes detected in the particulate organic matter fraction of surface water (SRF) and bottom of the surface mixed layer (BML).** The abundance of an epitope was determined from the antibody signal intensity and is expressed as relative to all samples for that epitope. Therefore, the abundance of an epitope can be compared across amples but not to that of epitopes from other antibodies. Specific polysaccharide structures that the molecular probes (depicted in parentheses) bind to are shown in the legend. FCSP = fucose-containing sulphated polysaccharide.

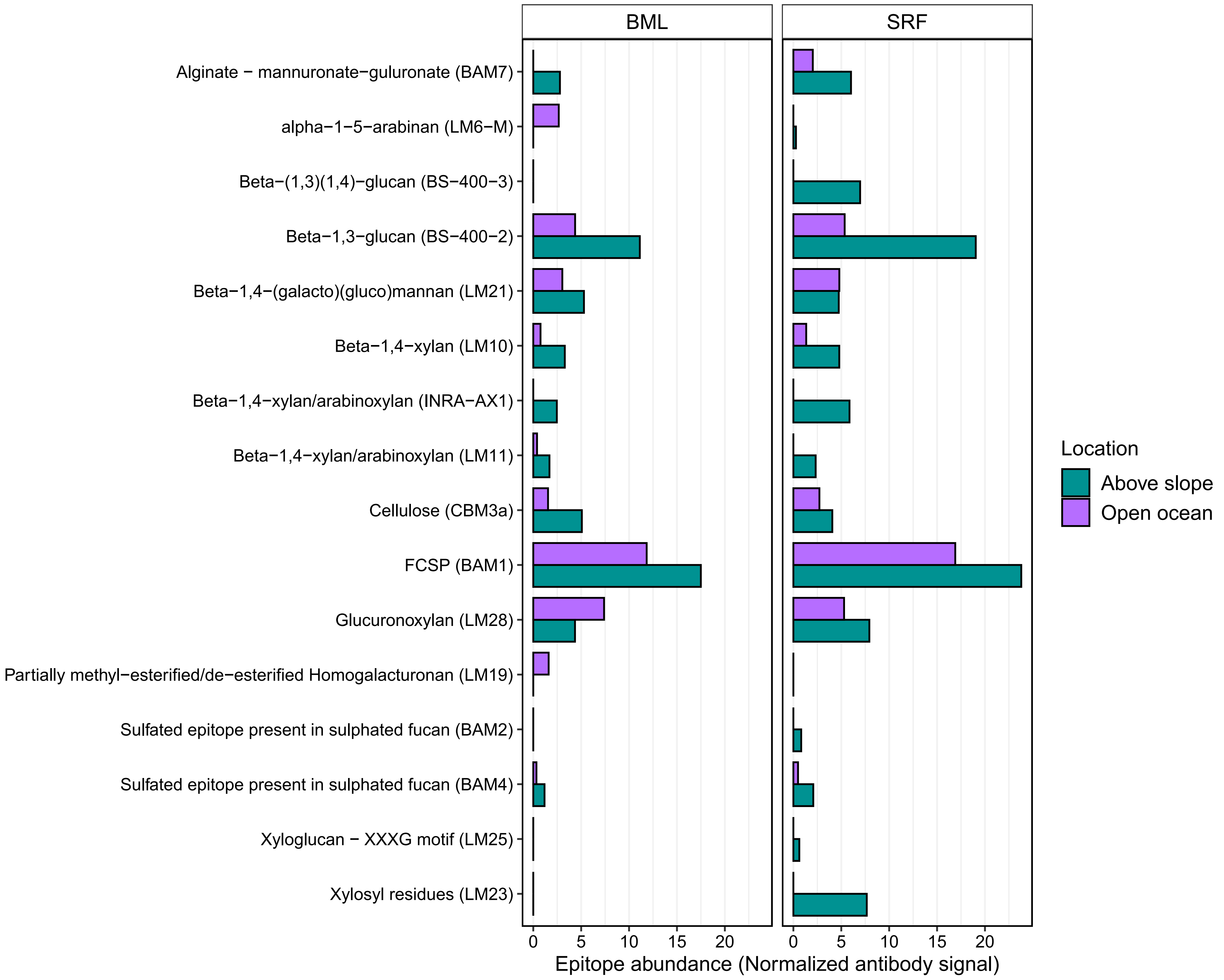


**Supplementary Figure S3. Mean abundance of polysaccharide epitopes in the particulate organic matter in surface water (SRF) and bottom of the surface mixed layer (BML) of open ocean and above continental slope locations.** The abundance of each epitope was derived from the signal intensity of antibody binding. The highest signal intensity was set to 100 and all other values normalized accordingly. The mean abundance was then taken for each epitope across above slope and open ocean samples. As a result, the abundance of different epitopes can not be directly compared but the abundance of the same epitope across locations can be compared.

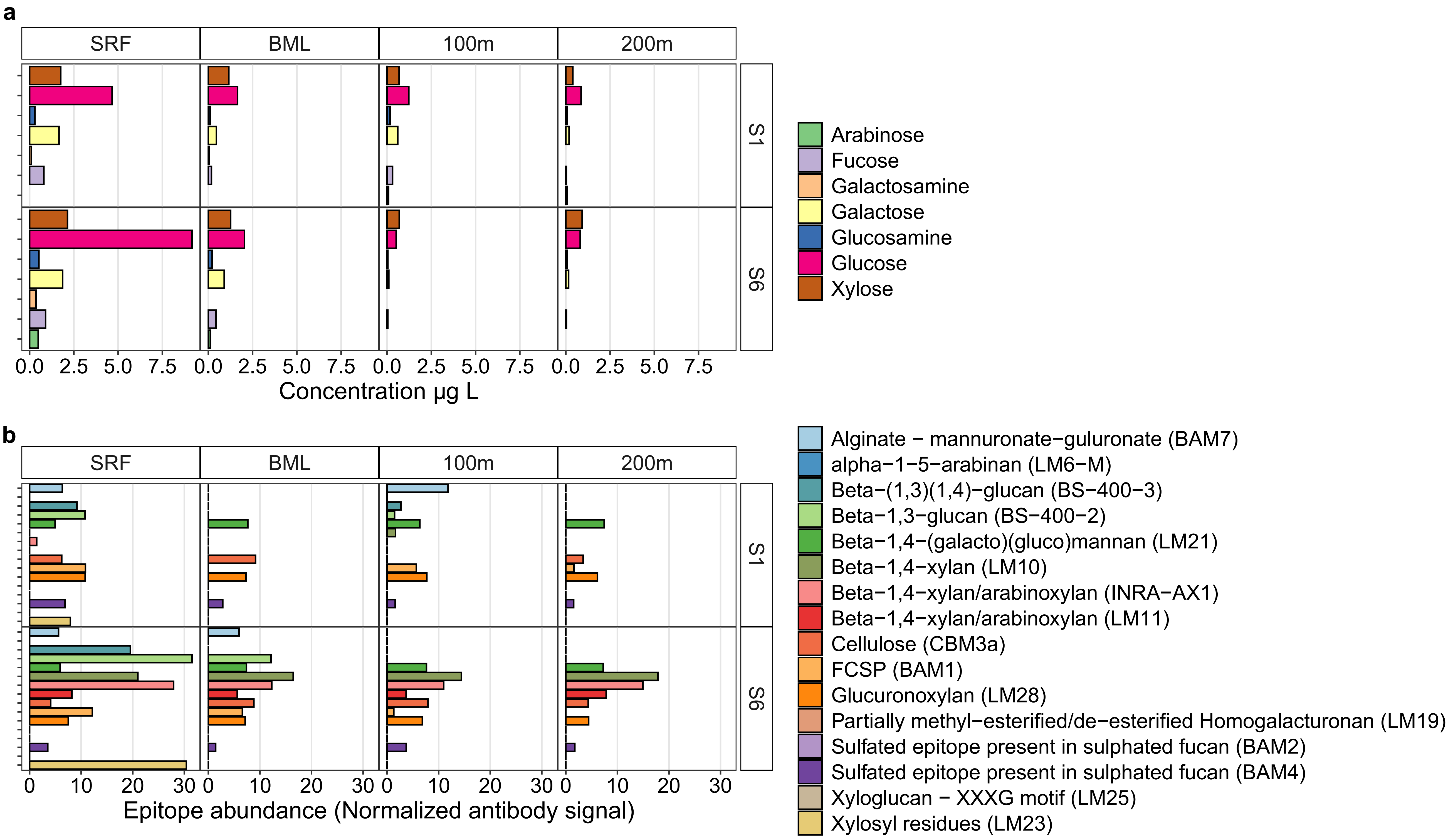


**Supplementary Figure S4. Monosaccharide and polysaccharide composition of the particulate organic matter fraction from surface to 200 m depth.** SRF = surface water, BML = bottom of surface mixed layer. The two stations represent the same location sampled over a two day period. Station S1 samples were collected 08:11 and station S6 samples at 16:30 (+24 h). Panel **b)** legend shows the polysaccharide structures that the corresponding monoclonal antibodies (depicted in parentheses) bind to. FCSP = fucose-containing sulphated polysaccharide.

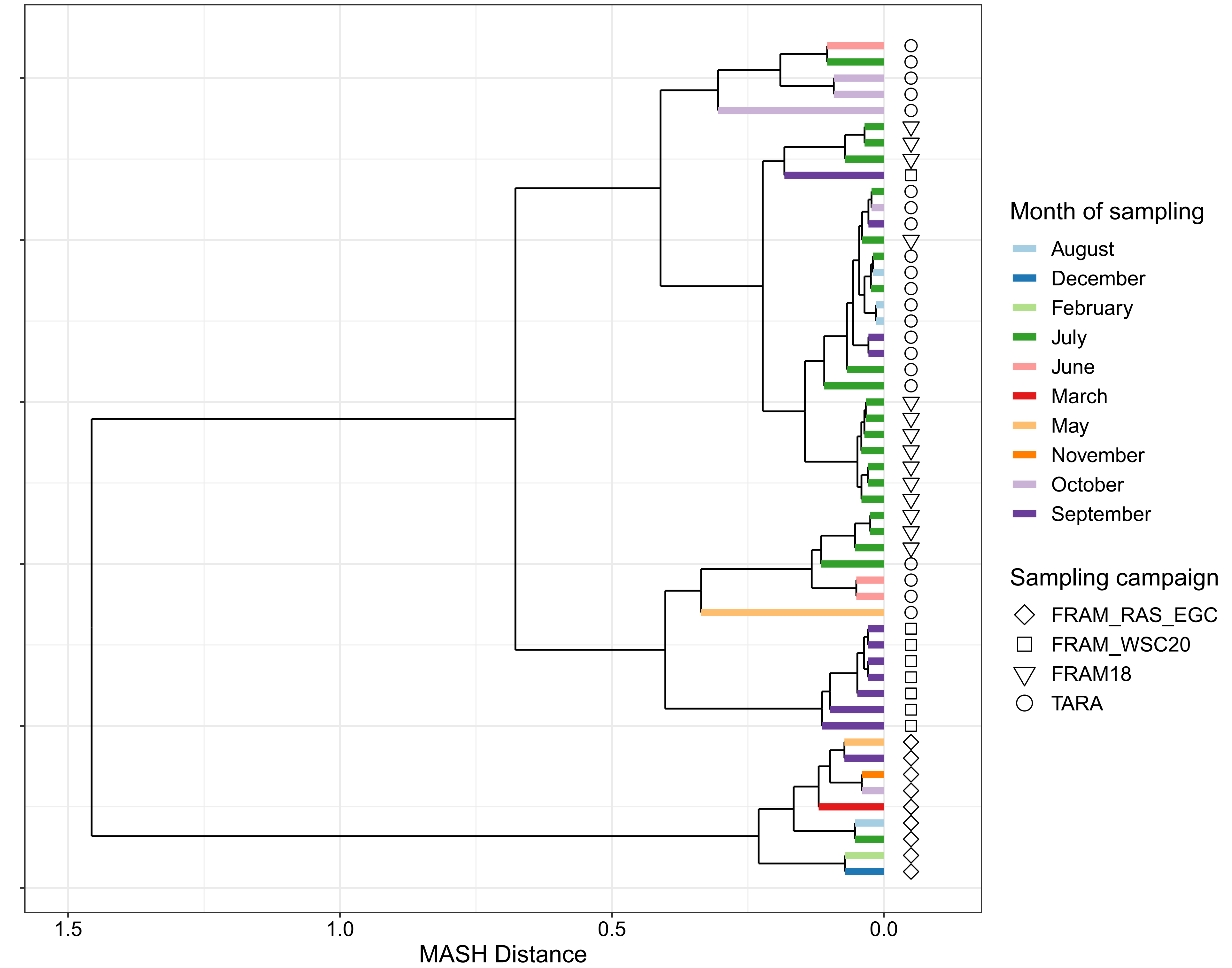


**Supplementary Figure S5. Hierarchical clustering of metagenomes from this study and those previously published from the Fram Strait and Arctic Ocean based on sequence composition.** Comparison of sequence composition between metagenomes was performed using MASH, with the resulting MASH distances being used as an input for hiearchical clustering using the average linkage algorithm. FRAM_WSC20 = samples from this study, FRAM18 = samples from PRJEB41592, FRAM_RAS_EGC = samples from PRJEB52171, TARA = samples from PRJEB9740.

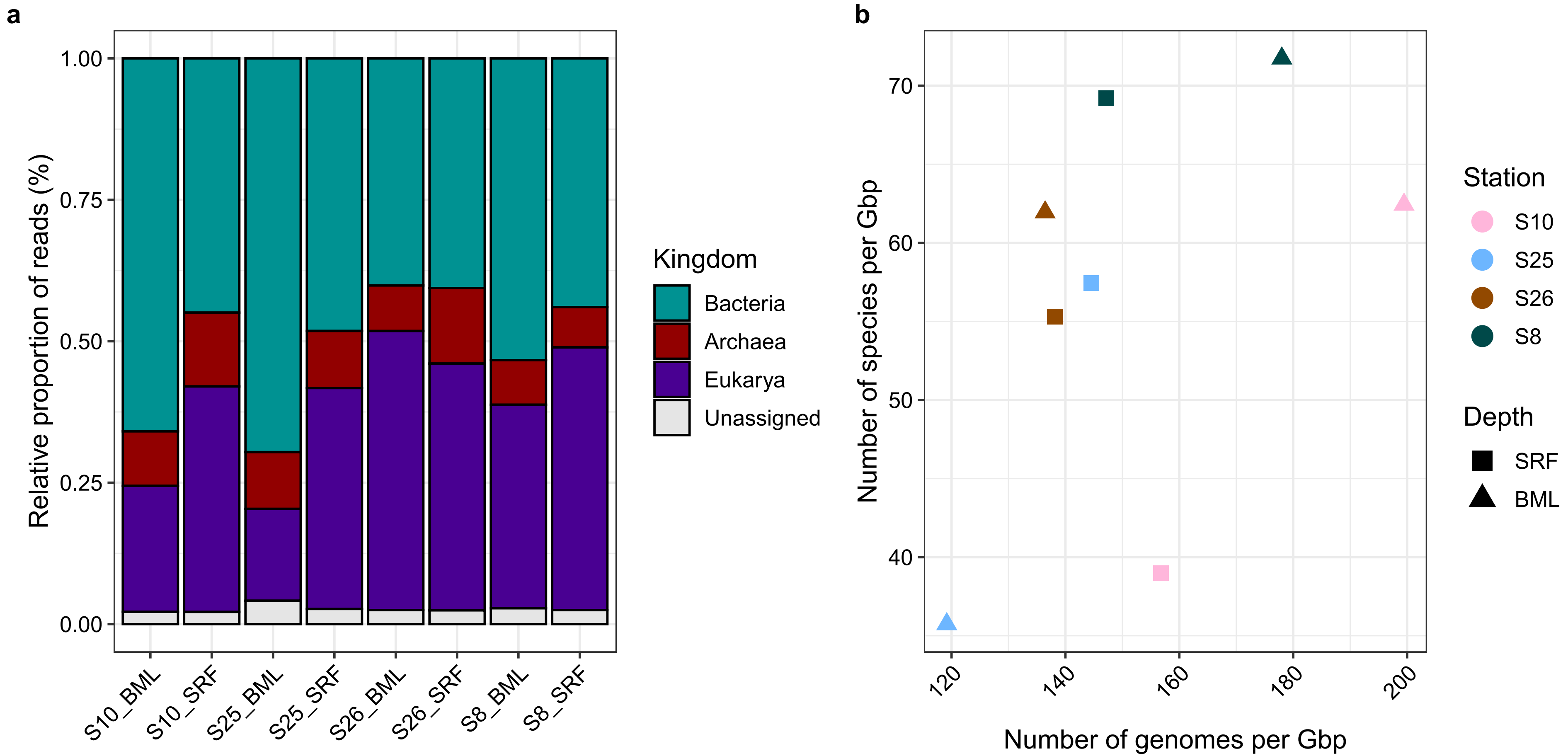


**Supplementary Figure S6. Composition, diversity and complexity of metagenomes. a)** Domain-level taxonomic composition of metagenomic reads. **b)** The number of microbial genomes and species captured in metagenomes derived from single-copy ribosomal protein gene sequencing depth and previously published species delineation thresholds.

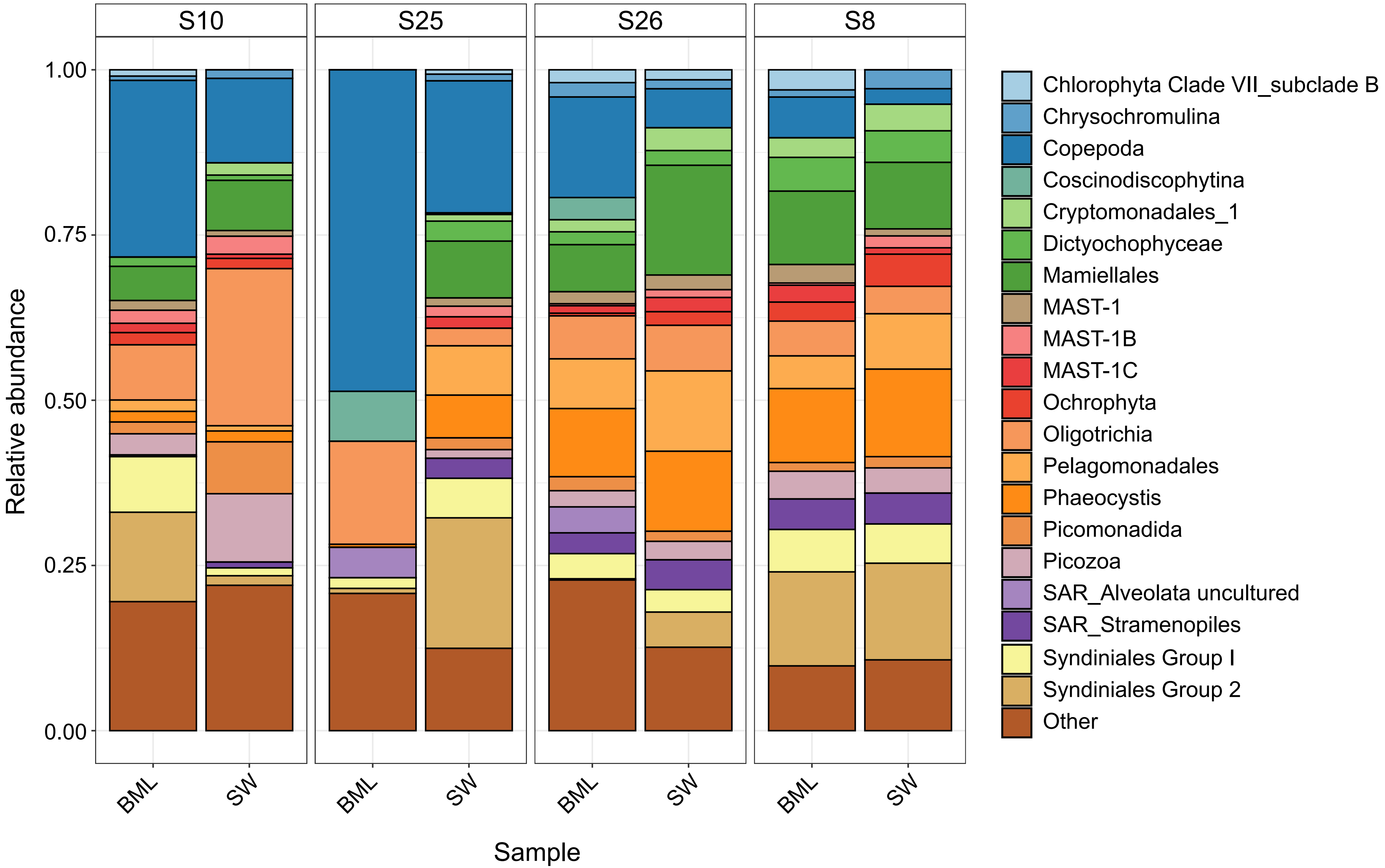


**Supplementary Figure S7. Composition of 18S rRNA gene sequences extracted from metagenomic reads.** The 18S rRNA gene sequences were extracted from the metagneomic reads using Barrnap and processed using DADA2 with the SILVA SSU Ref138.1 database for classification.

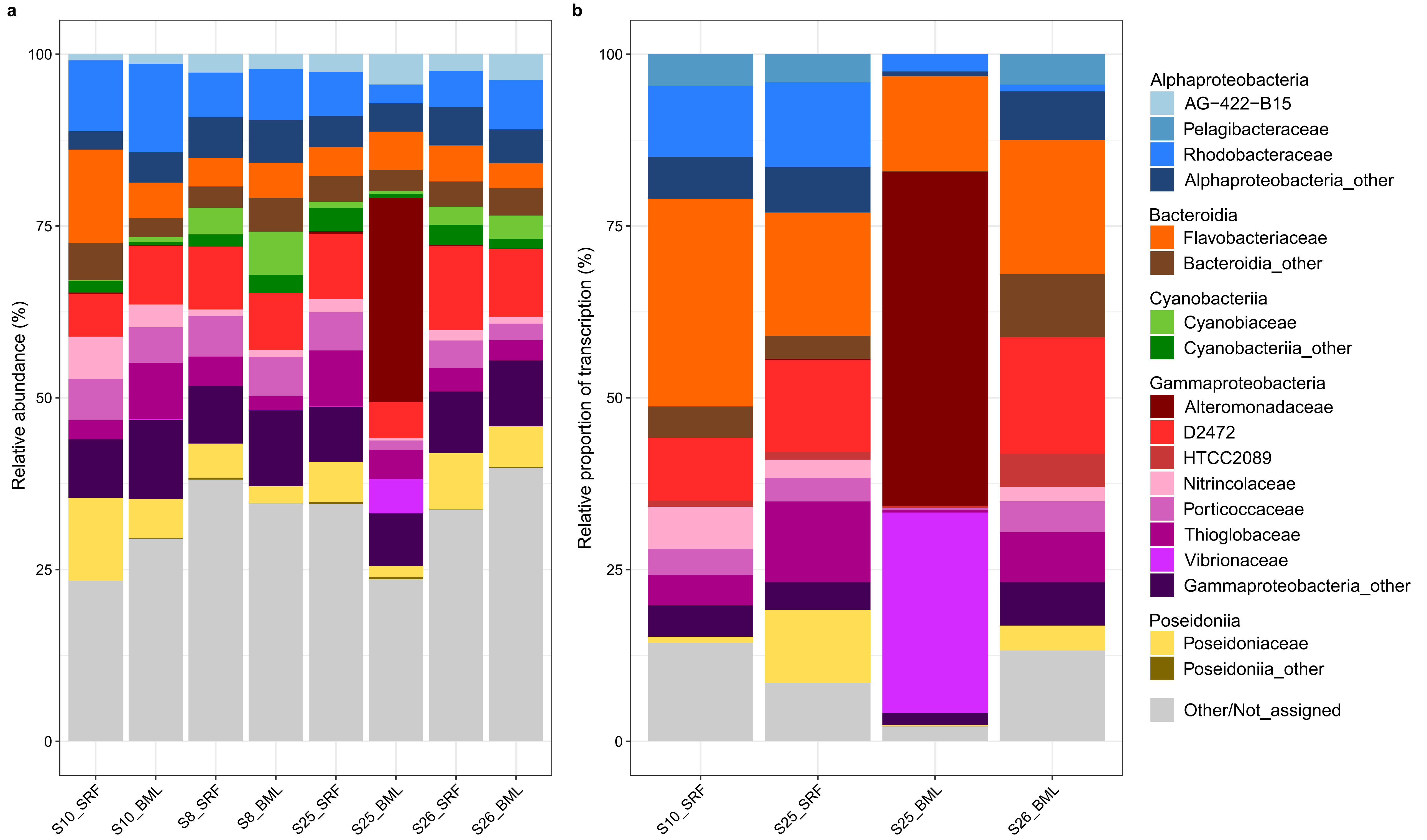


**Supplementary Figure S8. Family-level composition of microbial communities in the metagenomes and metatranscriptomes. a)** Relative abundance of families based on proportional sequencing depth of family-assigned ribosomal protein gene L6 compared to sample total sequencing depth of gene. **b)** Relative proportion of transcription of ribosomal protein gene L6 to sample total transcription of gene. Taxonomic classificaiton was derived from classifying the original HiFi reads, from which the genes were derived, against a GTDB-based protein database.

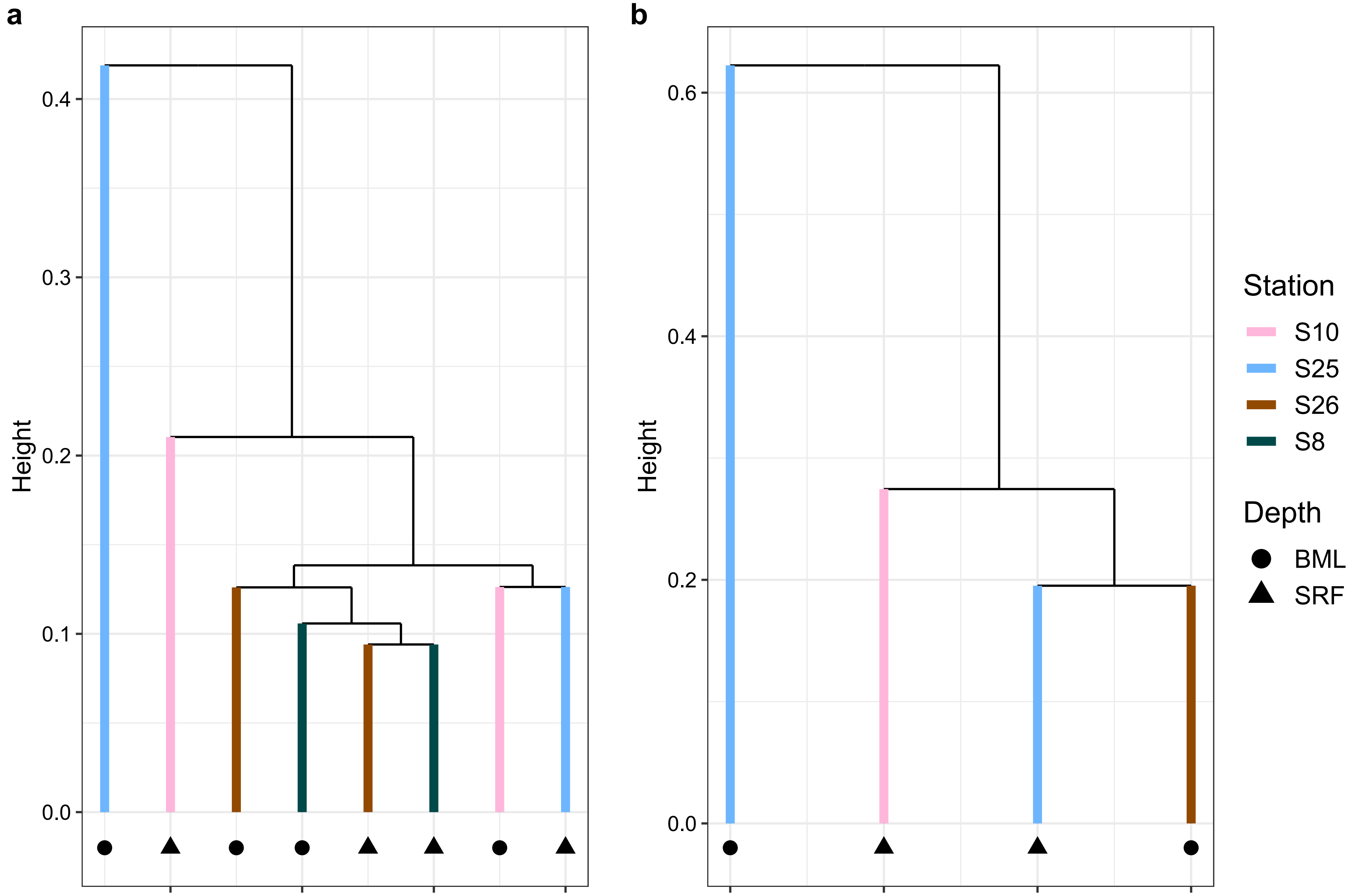


**Supplementary Figure S9. Dissimilarity in the CAZyme gene composition of metagenomes and metatranscriptomes.** The abundance **(a)** and transcription **(b)** of carbohydrate-active enzyme (CAZyme) genes was normalised by the number of microbial genomes present in each sample and subsequently converted to a Bray-Curtis dissimilarity matrix. The matrices were used for hierarchical clustering with the average linkage algorithm.

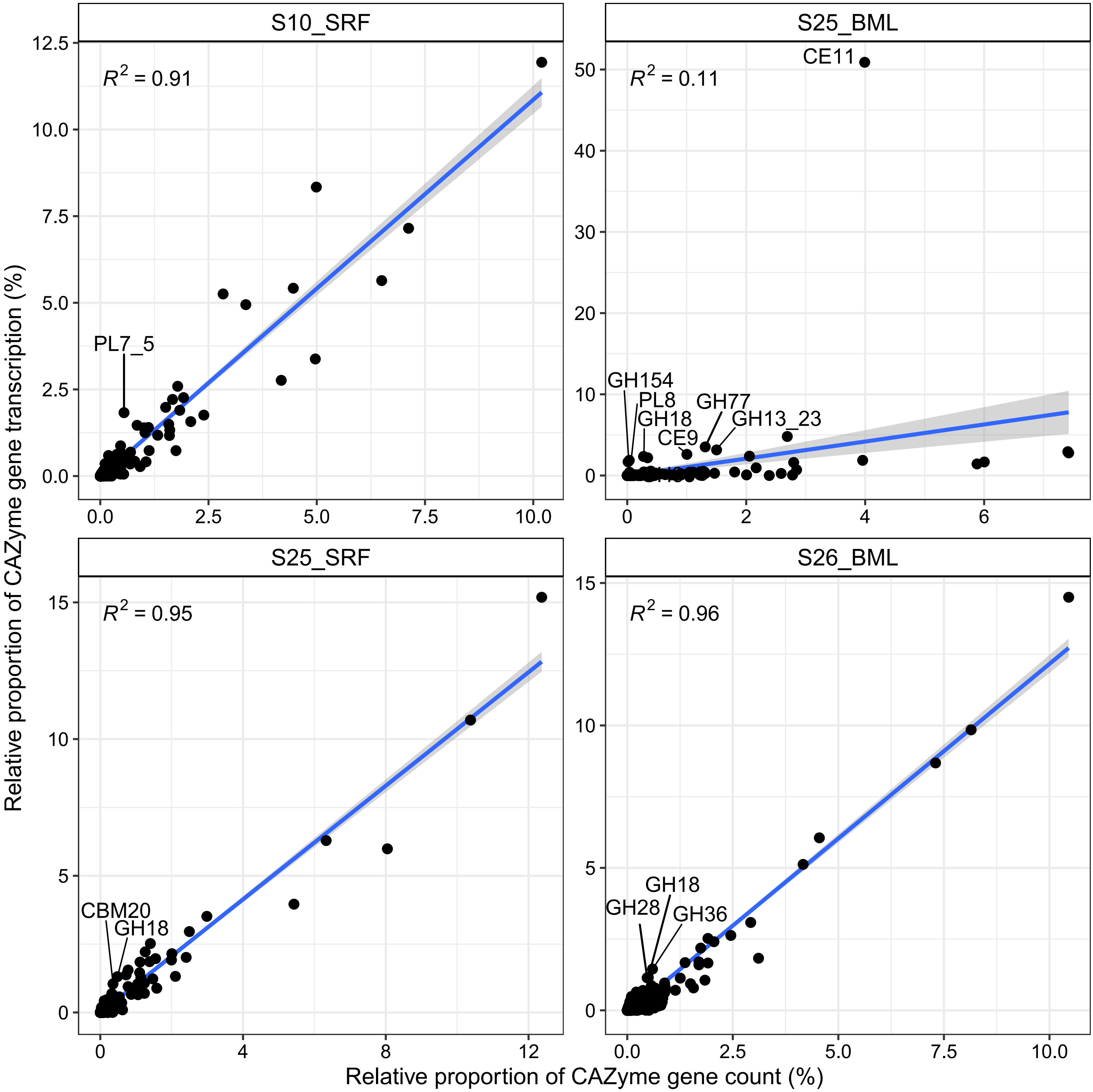


**Supplementary Figure S10. Linear relationship between CAZyme gene family abundance and transcription.** CAZyme gene counts and transcription (transcripts per million) values were summed at the gene family level before being normalised by the number of microbial genomes in each sample and then converted to relative proportions. CAZyme gene family points with labels are those where the relative proportion of transcription was more than double the relative proportion of abundance and with a relative transcription of >1%.

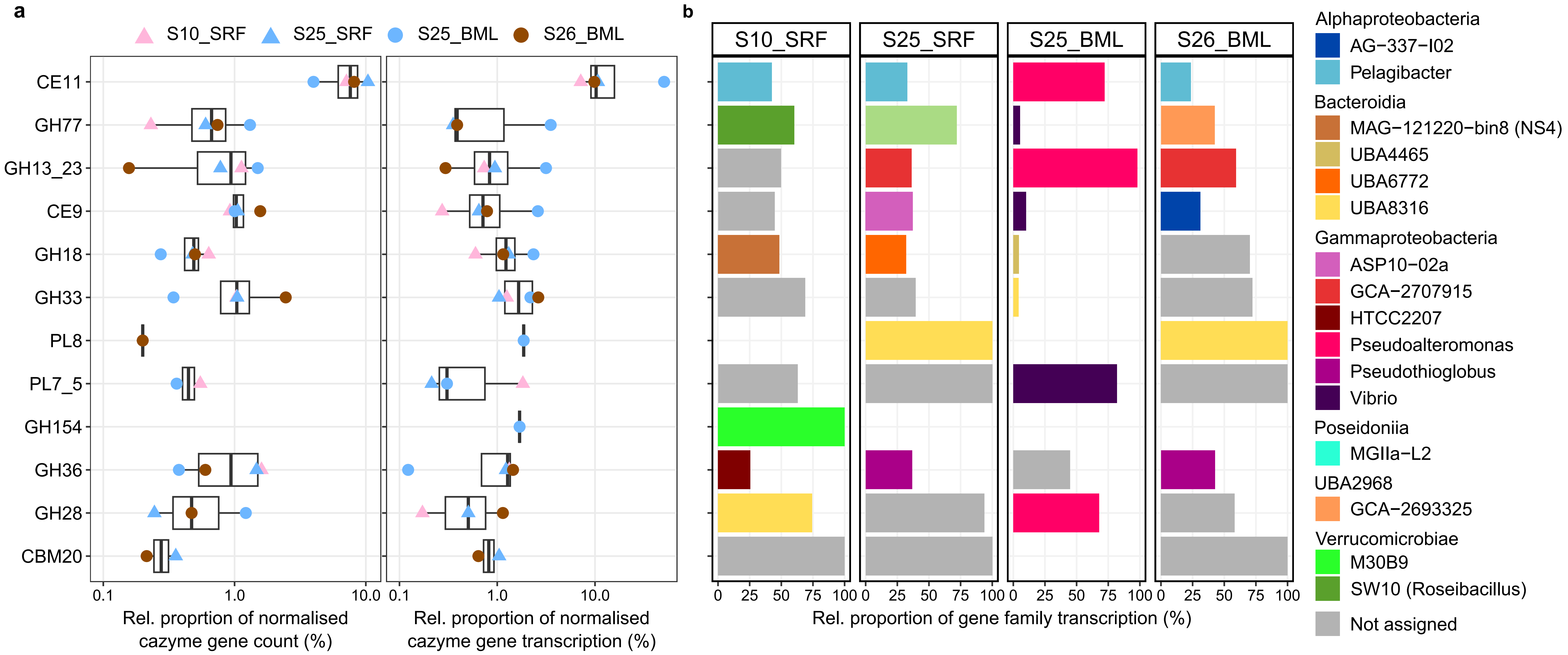


**Supplementary Table S11. Abundance, transcription and taxonomic information of the CAZyme gene families with a disproportionately high level of transcription compared to abundance. a)** The abundance and transcription of carbohydrate-active enzyme (CAZyme) genes was normalised by the number of microbial genomes per sample and then converted to relative proportions. The selected CAZyme gene families visualised are those that exhibited a twofold higher relative proportion of transcription than relative proportion of abundance in at least one sample. **b)** The genera that contributed the most to transcription of each gene family in each sample. Taxonomic classification of CAZyme genes was derived from the classification of the read from which it was derived.
