## Supplementary Methods for "Spatial heterogeneity in carbohydrates and their utilisation by microbes in the high North Atlantic"

*Monosaccharide and polysaccharide analysis*

The GF/F filters from all samples (ten stations and two depths) were cut into twelve equally-sized circular pieces, with a diameter of 11.2 mm. For monosaccharide analysis, two of the circular sections from each sample were subject to acid hydrolysis in glass ampules with 500 µl of 1 M HCl for 24 h at 100 °C. A 450 µl aliquot was then transferred to a centrifugal vacuum concentrator for drying, before resuspension in 150 µl of MilliQ water. The resuspended material was briefly vortexed, spun down and transferred to a high-performance liquid chromatography (HPLC) glass vial. The samples were analysed using high-performance anion exchange chromatography with pulsed amperometric detection along with monosaccharide standards, as described in [22]. Due to a problem with the detector neutral and amino monosaccharides could be detected but not the acidic sugars.

For carbohydrate microarray analysis, ten circular sections from each sample were subject to a sequential polysaccharide extraction protocol with 1) MilliQ water, 2) 0.2 M EDTA (pH 7.5) and 3) 4 M NaOH with 0.1% w/v $NaBH_4$. In each round of extraction, filter sections were placed in a 2 ml tube with 450 µl of the respective solvent, briefly vortexed and then incubated for 2 h at 60 °C in a tube shaker at 650 rpm. After incubation, the tubes were centrifuged at 6000 x g for 15 min at 15 °C. The supernatant was transferred to a new 1.5 ml tube. The next solvent was added to the pellet and the same process was repeated. The identification and semi-quantitative analysis of polysaccharide epitopes was performed using a microarray- and antibody-based approach, as described previously [22]. Briefly, polysaccharide extracts were loaded into wells of 384 microwell-plates and centrifuged at 3500 x *g* for 10 min at 15 °C. The contents of the wells were printed onto nitrocellulose membranes (0.45 µm pore size) using a microarray robot (Sprint, Arrayjet, Roslin, UK) under conditions of 20 °C and 50% humidity. Extracts were printed in quadruplicates, so that each extract was represented by 4 spots in the microarray. The printed arrays were blocked for 1 h in 1 x PBS with 5% (w/v) non-fat milk powder (MPBS). The MPBS was subsequently discarded and the microarrays were individually incubated for 2 h with one of 50 polysaccharide-specific monoclonal antibodies diluted in MPBS (see the antibodies and dilution factors in Supplementary Table S2). After incubation, the arrays were washed in PBS and incubated for 2 h in anti-rat, anti-mouse or anti-His-tag secondary antibodies conjugated to alkaline phosphatase diluted 1:5000, 1:5000 and 1:1500 in MPBS, respectively. Arrays were thoroughly washed in PBS, followed by deionised water and developed in a solution containing 5-bromo-4-chloro-3-indolylphosphate and nitro blue tetrazolium in alkaline phosphatase buffer (100 mM NaCl, 5 mM $MgCl_2$, 100 mM Tris-HCl, pH 9.5). Developed arrays were scanned and the binding of each probe against each spotted sample was quantified using Array-Pro Analyzer 6.3 (Media Cybernetics). Mean antibody signal binding intensity (n = 4) was then determined for each extract. The highest mean value of the dataset (which corresponded to a polysaccharide standard control) was set to 100 and all other values were normalized accordingly.
